# Environment Modulates Protein Heterogeneity Through Transcriptional and Translational Stop Codon Miscoding

**DOI:** 10.1101/2023.02.01.526636

**Authors:** Maria Luisa Romero Romero, Anastasiia Kirilenko, Jonas Poehls, Doris Richter, Tobias Jumel, Anna Shevchenko, Agnes Toth-Petroczy

## Abstract

Stop codon miscoding events give rise to longer proteins, which may alter the protein’s function and thereby generate short-lasting phenotypic variability from a single gene.

In order to systematically assess the frequency and origin of stop codon miscoding events, we designed a library of reporters. We introduced premature stop codons into mScarlet that enabled high-throughput quantification of protein synthesis termination errors in *E*.*coli* using fluorescent microscopy. We found that under stress conditions, stop codon miscoding may occur with a rate as high as 80%, depending on the nucleotide context, suggesting that evolution frequently samples stop codon miscoding events. The analysis of selected reporters by mass spectrometry and RNA-seq showed that not only translation but also transcription errors contribute to stop codon miscoding. The RNA polymerase is more likely to misincorporate a nucleotide at premature stop codons. Proteome-wide detection of stop codon miscoding by mass spectrometry revealed that temperature regulates the expression of cryptic sequences generated by stop codon miscoding in *E*.*coli*.

Overall, our findings suggest that the environment influences the accuracy of protein production, which increases protein heterogeneity when the organisms need to adapt to new conditions.

## INTRODUCTION

Protein synthesis termination is optimized to have high fidelity, yet it is not perfect. Within all life forms on Earth, the translation end is signaled by stop codons and catalysed by release factors^1–3^. The process has evolved to rewrite genomic information into canonical-size proteins accurately^4^. However, stop codon miscoding (SCM) may occur either by a transcription error, when the RNA polymerase misincorporates a nucleotide and eliminates the stop codon, or by a translation error when the ribosome misincorporates a tRNA at the stop codon (also called stop codon readthrough and nonsense suppression^5,6^). These errors result in protein variants with extended C termini^7–9^, generating a heterogeneous protein-length population from a single gene.

Proteome diversification arising from SCM can potentially generate phenotypic variability, shaping the cell’s fate^10^. For instance, infidelity in protein synthesis termination can be adaptive and functional^7,9,11–16^ or non-adaptive^17^, and it sometimes leads to fitness decrease^18^. Further, slippery sequences upstream the stop codon that cause the ribosome to slip and thereby lead to stop codon readthrough are often functional^19,20^. Transcription and translation errors may generate short-lasting phenotypic variability on a physiological time scale, faster than genomic mutations. Thus, SCM events may facilitate rapid adaptation to sudden environmental changes. These tremendous evolutionary implications reveal the need to study the rules that dictate inaccuracy of protein synthesis termination under diverse living conditions.

Few studies have highlighted the relevance of stop codon miscoding under diverse environmental conditions. Both carbon starvation^21^ and excess glucose promote the readthrough of TGA in *E. coli* by lowering the pH^22^. However, these studies provided information for a given stop codon and genetic context, detecting a maximum of 14% TGA readthrough when cells were grown in LB supplemented with lactose^22^. Further, low growth temperature dramatically increased ribosomal frameshift rate in *Bacillus subtilis*^23^. Nevertheless the impact of temperature on SCM is unknown.

Here we systematically examined the inaccuracy of protein synthesis termination of all stop codons in a variety of genetic contexts under different temperatures and nutrient conditions. We specifically asked: i) how frequent errors are in protein synthesis termination, ii) whether non-optimum environmental conditions modulate the chances for evolution to encounter these events; and iii) what their origin is, translation or transcription inaccuracy. We designed a library of reporters that allowed for high-throughput quantification of protein synthesis termination errors in *E. coli* using fluorescent microscopy. We targeted 43 arbitrary positions along the mScarlet sequence, and, in each position, we mutated the wild-type codon to each of the three stop codons. Thus, only upon errors that result in skipping protein synthesis termination, full-length mScarlet will be synthesized.

We confirmed that protein termination accuracy depends on the identity and genetic context of the stop codon. We thus proposed a set of simple rules to predict hotspots in the protein sequence that are error-prone for protein synthesis termination. We further showed that environmental stress conditions such as low temperature and nutrient depletion increase the miscoding rates of all stop codons. Accordingly, the opal stop codon TGA, present in 29% of the *E. coli* proteome, fails to terminate the protein synthesis at certain positions at a rate of up to 80% under stress conditions at a given nucleotide context. RNA-seq and mass spectrometry experiments of selected reporters revealed that protein synthesis mis-termination is not only due to ribosomal readthrough, but RNA polymerase errors also contribute to SCM. We found that the RNA polymerase is more likely to misincorporate a nucleotide at premature stop codons. Finally, mass spectrometry analysis of *E. coli* proteome provided evidence of cryptic sequences revealed by SCM and validated our rules to predict SCM in a more natural context. Overall, our findings suggest a cross-talk between the environment and the biological flux of information that increases protein heterogeneity when organisms need to adapt to new conditions.

## RESULTS

### Visualizing and quantifying stop codon readthrough events in *E. coli*

To monitor inaccuracy in protein synthesis termination, inspired by the pioneering work of R.F. Rosenberger and G.Foskett^24^, we designed a fluorescence reporter whereby stop codon miscoding can be visualized and quantified in *E. coli*. The strategy dwells on introducing a premature stop codon into an mScarlet allele. Thus, only upon stop codon miscoding, full-length and, therefore, functional mScarlet will be synthesized (Fig 1A).

**Figure 1.**
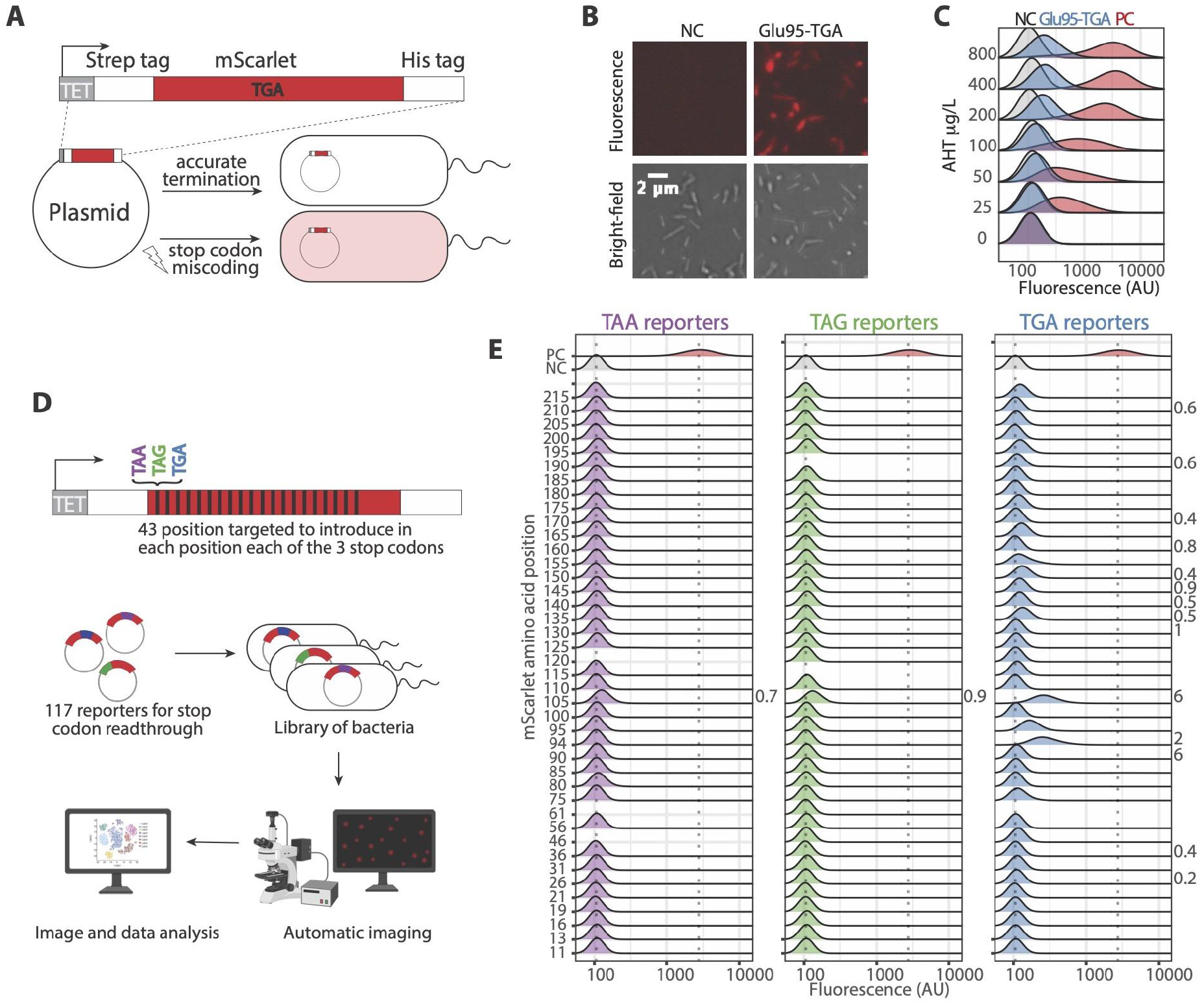
Visualization and quantification of stop codon miscoding events in *E. coli* using fluorescence reporters. **A)** To study stop codon miscoding (SCM), we designed a fluorescence reporter introducing a premature stop codon into an mScarlet allele. **B)** We detected full-length mScarlet and, therefore, SCM events, in cells transformed with the reporter. **C)** Titrating the expression of the reporter increases the detection of SCM events. **D)** We designed a library of reporters for SCM mutating to TAA (purple), TAG (green), and TGA (blue), 41 randomly selected codons along the mScarlet sequence. We studied the library of reporters in a high throughput fashion. **E)** Fluorescence distributions displayed by the *E. coli* cells, transformed with each library’s reporters and grown at 37°C in rich media. The propensity of SCM, calculated as the percent of the median fluorescence compared with the positive control, is shown for the distributions with a median fluorescence higher than the negative control. There are hotspots in the protein sequence prone to SCM. The identity of the stop codon influenced the likelihood of stop codon readthrough with the trend TGA > TAG > TAA.

We first introduced a TGA stop codon in the randomly-selected position 95 of an mScarlet flanked by two tags, Strep-tag at the N-terminal and His-tag at the C-terminal. These tags provide, on the one hand, a way to purify expressed fragments. On the other hand, the His-tag serves as an orthologous method to detect stop codon readthrough events by Western blot. To tightly regulate the expression of the reporter, we used a low copy number vector and an inducible promoter.

We transformed a wild-type *E. coli* strain with the Gly95-TGA reporter. Then we grew the transformed cells until stationary phase under optimal conditions, testing the expression of the Gly95-TGA reporter with titrated concentrations of the inducer. We used as negative control (NC) cells transformed with the empty vector and as positive control (PC) cells expressing wild-type mScarlet (Fig 1B and C). While no signal was detected in the negative control, an increased fluorescent signal was detected in the cells expressing the Gly95-TGA reporter and in the positive control, the higher the inducer’s concentration was. These results suggest that functional mScarlet was expressed in the cells carrying the Gly95-TGA reporter, meaning stop codon miscoding occurred. Assuming that the fluorescence properties of the SCM variants were not altered, we can quantify the error rate by calculating the relative fluorescence signal as the percentage of the median compared with the PC median. We found that stop codon readthrough is not a rare event; protein synthesis termination fails for 2.1 % of the expressed Gly95-TGA reporters (Fig 1C).

Since we compare cells with orders of magnitude difference in fluorescence intensity, we confirmed that the fluorescence measurements were within the dynamic range of the instrument, and the relationship between mScarlet concentration and fluorescent intensity is linear (Fig S1). Therefore, relative fluorescence is a valid approximation for the relative protein abundance, which allows us to quantify the stop codon miscoding error rate.

### Stop codon miscoding events are frequent

We, and others^23,25–28^, showed proof-of-principle that stop codon miscoding occurs at the opal stop codon, TGA and can be visualized and quantified with a fluorescent reporter. Next, we extended the strategy to study how the identity of the stop codon, the sequence, and the structural context regulate the inaccuracy in protein synthesis termination. To address these questions in a high-throughput fashion, we designed a library of reporters for stop codon miscoding. We randomly targeted 43 codons along the mScarlet sequence and mutated them to each of the three stop codons. We confirmed with sequencing that the final library consists of 117 reporters, 38 with TAA, 39 with TAG, and 40 with TGA stop codon. We transformed *E. coli* with the reporters individually and grew them, inducing the reporter expression under normal conditions in a 384-well plate. Next, we automatically imaged the cells in stationary phase to determine fluorescence (Fig 1D, see methods for detailed description).

Several reporters displayed high fluorescence output, meaning full-length mScarlet was expressed (Fig 1E). This is a rather unexpected result because the premature stop codons were introduced randomly along the mScarlet sequence, without consideration of the genome context that may promote stop codon miscoding. Still, the fluorescence distribution of the cells carrying a reporter with a premature stop codon at position 105 displayed a median fluorescence varying from 0.7 to 6% relative to the median fluorescence of the positive control (Fig 1E).

The identity of the stop codon seemed to be pivotal in regulating the stop codon miscoding propensity. Being TGA the most error-prone stop codon: 14 out of the 40 reporters carrying a premature TGA displayed a median fluorescence above the threshold defined as the mean plus two standard deviations of the fluorescence signal of the NC. Further, we found that the identity of the stop codon influenced the likelihood of stop codon miscoding and identified the trend TGA > TAG > TAA when the premature stop codon was in position 105 (Fig 1E), in agreement with previous observations in bacteria^29^, yeast^30^ and mammals^15,31^. Surprisingly, the least efficient stop codon, TGA, is present in 29% of the *E. coli* proteome. However, TAG, which is less prone to be miscoded, is only present in 8% of the *E. coli* proteome^32^.

In summary, our results suggest that stop codon miscoding events are more frequent than previously thought.

### Non-optimal growth temperatures and nutrient scarcity promote stop codon miscoding

Stop codon miscoding (SCM) events can generate short-lasting phenotypic variability faster than genomic mutations. Thus, SCM events may facilitate rapid evolution to sudden environmental changes. However, up to now, little attention has been paid to address how environmental conditions affect SCM. It is known that the readthrough of stop codons in *E. coli* depends on the growth media^21,22^. However, how temperature influences SCM remains unknown. To examine this question, we first screened our previously described library of reporters in wild-type *E. coli* grown at different temperatures (Fig 2A and S2).

**Figure 2.**
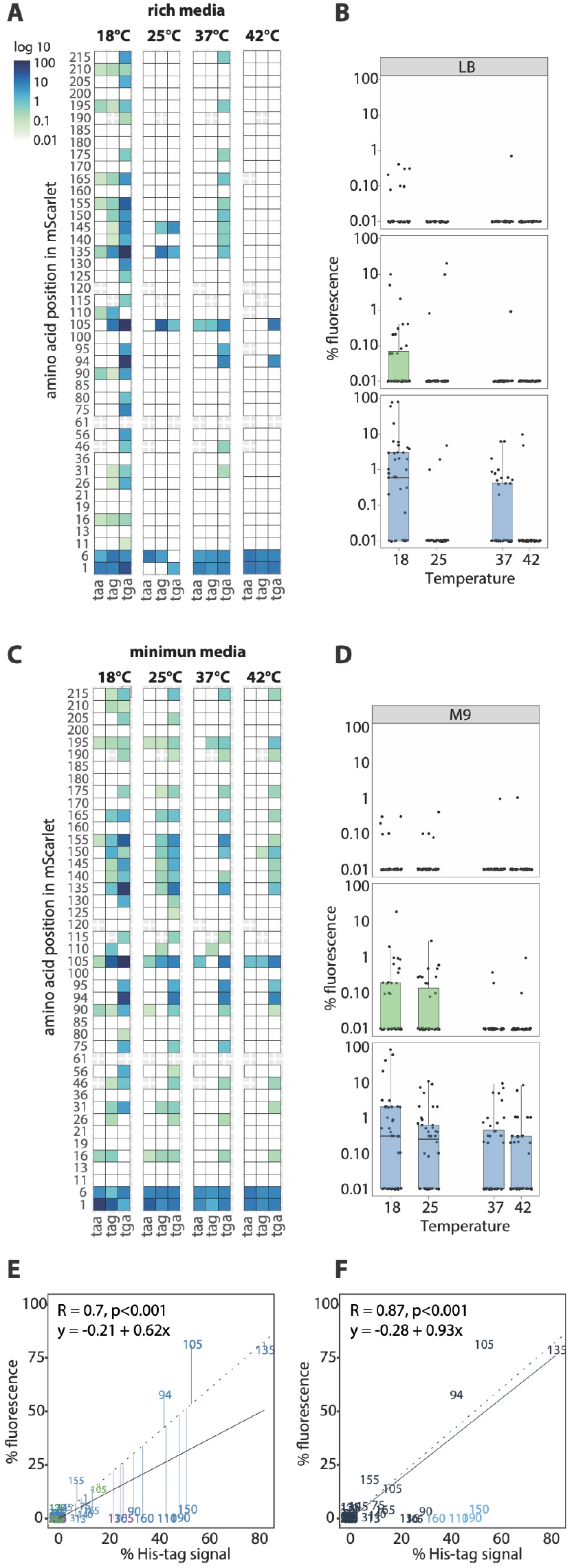
Non-optimal growth temperatures and nutrient scarcity promote stop codon miscoding. **A)** In cells grown in rich media, the stop codon miscoding (SCM) levels vary considerably as a function of the temperature. At 18°C more reporters displayed SCM events and at a higher rate. **B)** When cells were grown in rich media, at all tested temperatures, TAA was the most accurate codon terminating the protein synthesis and TGA the least. **C)** Nutrient scarcity promoted SCM. In cells grown in minimum media, SCM levels increased while lowering the growth temperature. **D)** When cells are grown in minimum media, at all tested temperatures, TAA is the most accurate codon terminating the protein synthesis and TGA the least. **E)** C-terminal His-tag expression correlates with fluorescence signal. Only four out of 114 reporters underestimate the rate of SCM events likely due to loss of fluorescence (“dark reporters”). **F)** Excluding the four dark reporters, the linear fit between the percentage of His-tag and fluorescence signal presents a slope of 0.93, close to the ideal value of 1.

A closer inspection of the temperature-effect scan revealed that there are hotspots in the sequence prone to SCM. We define hotspot positions, where the premature stop codon is miscoded in more than 50% of the tested conditions, for example, positions 105 and 135 (Fig 2A). These hotspots seem to depend on the sequence context of the premature stop codon since, regardless of the stop codon identity, they are likely to err the protein synthesis termination at different temperatures.

Within all temperatures, TAA is least likely to be miscoded while TGA is most likely to be miscoded, in agreement with previous observations^31,33,34^. At each temperature, more TGA reporters display stop codon miscoding than TAG and TAA (Fig 2A, 2B, and Fig S2). Also, at the hotspot positions, the stop codon termination error rates follow the same trend: TGA > TAG > TAA (Fig 2C).

The SCM rates vary considerably as a function of the temperature. Interestingly, at lower temperatures, more reporters display stop codon miscoding events (Fig 2A, FigS2, and Fig S3AB). At the hotspot positions, the protein synthesis termination seems more accurate at optimal growth temperature, 37°C (Fig S2). However, at low temperatures (18°C), when TGA is at position 135, the SCM is surprisingly frequent: the SCM rate is 80%. In order to test if the observed temperature effect is an artifact due to the unfolding of mScarlet at high temperatures, we assayed the stability of mScarlet, showing that mScarlet has an apparent melting temperature (T_m_) above 80°C (Fig S3C).

To examine how nutrient depletion modulates protein synthesis termination accuracy, we screened our library of reporters in wild-type *E. coli* grown in minimal-medium, M9, at different temperatures (Fig 2D and S2). The most surprising aspect of these data is how inaccurate protein synthesis termination is when nutrients are limited. Indeed, the number of hotspots prone to stop codon readthrough increases to 10 (position 16, 90, 105, 135, 140, 150, 155, 165, and 195; Fig 2D). Besides the higher level of inaccuracy, the stop codon identity (Fig 2E, S2, S3AB) and the temperature effect (Fig 2F and S2) appear to play a similar role when growing *E. coli* in minimal-media than in rich-media. I.e., the stop codon miscoding level follows the TGA > TAG > TAA trend and is less likely at optimal growth temperatures.

To further explore the key nutrients essential to keep an accurate protein synthesis termination, we first supplemented the minimum media with a higher carbon source concentration (1.6% glycerol, Fig S4A) and, secondly, with a higher casamino acid concentration (0.4% casamino acid, Fig S4B). In both cases, the protein synthesis termination accuracy increases. I.e., when cells were grown in not supplemented minimum media, 18 reporters with TGA displayed a median fluorescence above the threshold (defined as the mean plus two standard deviations of the fluorescence signal of the NC). However, when supplementing with higher glycerol and casamino acid concentration, only 6 and 8 reporters, respectively, presented a median fluorescence above the threshold. This suggested that both nutrients, the carbon source, and the casamino acids, are essential to prevent SCM events.

Global inspection of these results suggests that environmental stress conditions, such as non-optimal growth temperature or nutrient scarcity, promote stop codon miscoding and, thus, protein heterogeneity. Since sudden environmental changes can modulate protein heterogeneity in a rapid and short-lasting fashion, we hypothesize that *E. coli* can use stop codon miscoding to quickly adapt to sudden and short-lasting stress conditions.

### C-terminal His-tag detection as an orthologous method to quantify stop codon miscoding events correlates with fluorescence signal

We linked the fluorescence signal of mScarlet with stop codon miscoding events. However, it could be that the miscoding of a stop codon disrupts the mScarlet tertiary or secondary structure resulting in full-length yet non-functional (i.e., non-fluorescent or dark) mScarlet or in mScarlet with potentially enhanced fluorescence. We used an orthologous method to detect SCM events and evaluated the limitations of our fluorescent reporters’ library. The reporters include a His-tag at the C-terminal (Fig 1A and 1D), and thus, only upon SCM, His-tag is synthesized. We used western blotting via anti-His tag antibodies as an orthologous method to detect SCM events.

We detected and quantified the His-tag expression of all the reporters expressed in wild-type *E. coli* grown at 18°C in LB media (Fig S5). We then correlated the percentage of the His-tag expression with the percentage of the fluorescence signal, using the positive control as the reference. Overall, there is a significant correlation between His-tag expression and fluorescence signal (Fig 2E). However, few reporters present a high His-tag expression but do not have fluorescence (dark reporters) (Fig 2F). We considered a reporter dark when the discordance between His-tag and fluorescence signal is over 30%. With this criterion, four reporters out of 114 were considered dark, and we excluded them for later analyses regarding the influence of the sequence context on SCM events. Without dark reporters, the linear fit between the percentage of His-tag and fluorescence signal presents a slope of 0.93, close to the ideal value of 1 (Fig 2F). Only one reporter (Ala-105-tga) presented a higher SCM error rate when assessing the fluorescence (80%) than when assessing the His-tag expression (53%). To rule out that this discordance is caused by changes in the mScarlet structure due to amino acid replacement at the stop codon site that enhances the mScarlet fluorescence, further experiments must be done.

Overall, the C-terminal His-tag analysis helped us to identify and deal with a limitation of our reporters’ design. The identified dark reporters underestimate the rate of SCM events in just four out of 114 reporters, proving that our reporter library is a valid method to study errors in protein synthesis termination.

### High G and low T content downstream of the stop codon increases the likelihood of stop coding miscoding

Our results show that there are hotspots in the sequence prone to SCM events. Since we targeted 43 positions in the mScarlet sequence, we can systematically compare the influence of the genetic context on SCM. We analysed the nucleotide occurrences surrounding the premature stop codon to see if they correlate with protein synthesis termination errors.

We derived a score to estimate how likely an SCM event is at a given position. Then, we binned the scored values and correlated them with the occurrence of each nucleotide in a 5-nt window upstream and downstream of the premature stop codon. The results show a significant positive correlation for G and negative for T content downstream of the premature stop codon and the likelihood of SCM (Fig 3A), contrary to previous observations in mammalian cells where the nucleotide at the +4 position increased readthrough in the following order C > U > A > G^31^. However, a non-significant correlation was found upstream of the premature stop codon (Fig 3B) (see Method section for further details).

**Figure 3.**
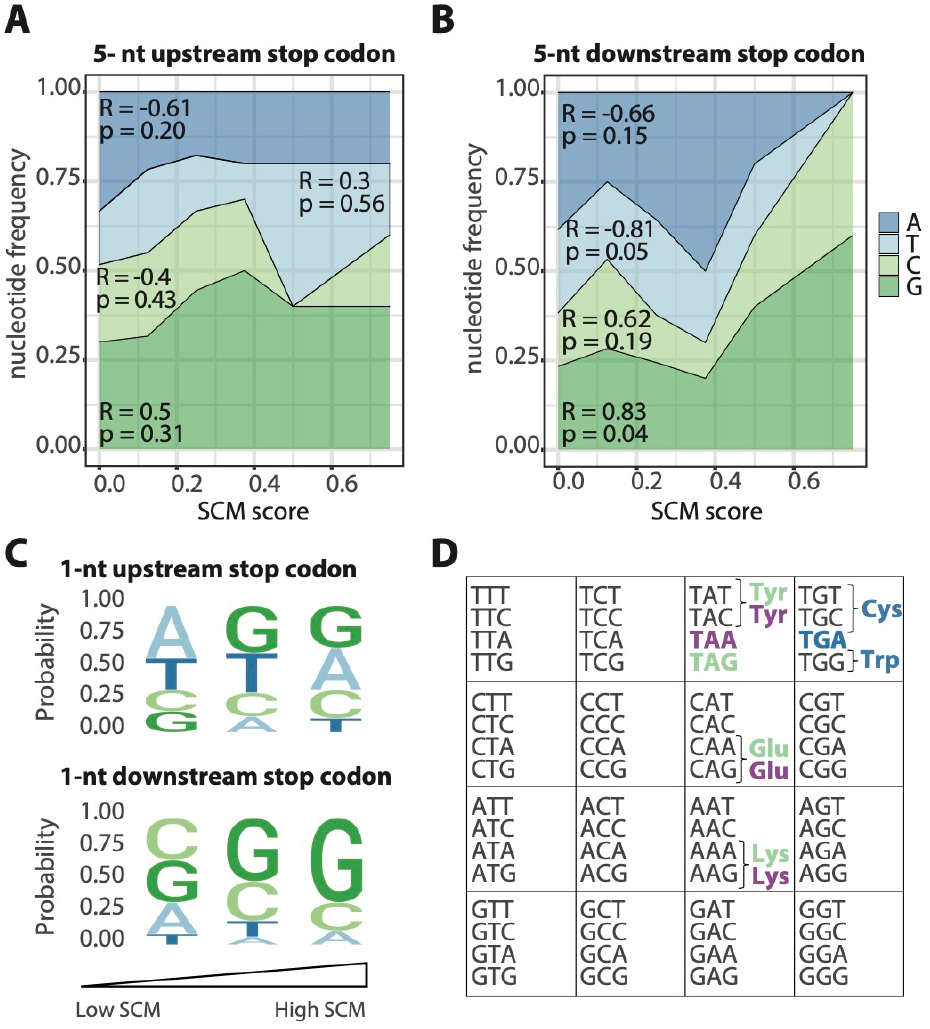
The nucleotide sequence downstream the premature stop codon influences the probability of stop codon miscoding. **A)** The identity of the nucleotides in a 5-amino acid window upstream the stop codon does not influence the probability of SCM. **B)** High G content in a 5-amino acid window downstream the stop codon increases the probability of SCM while T decreases it. **C)** Reporters were binned into three categories based on the SCM rates (see Methods) and the frequency of the nucleotides were calculated 1-nt upstream and downstream of the stop codons. The identity of the base immediately before the stop codon does not impact the probability of SCM. The identity of the base immediately after the stop codon impacts the probability of SCM, T increases, and G decreases the protein synthesis termination efficiency. **D) Stop codon miscoding events primarily occur due to amino acid misincorporations identified by mass spectrometry**. TGA is almost always replaced by tryptophan and, in lower frequency, by cysteine. TAG and TGA are replaced by various amino acids, such as glutamine, lysine, and tryptophan with similar frequency. Misincorporation of a few other amino acids at the stop codon was clearly a minor process (Table S1). Stop codon miscoding, SCM.

Since numerous reports have shown that the nucleotide immediately following the stop codon influences SCM in eukaryotes^31,35^ and prokaryotes^36^, we also studied the influence of the identity of the base following the stop codon on SCM. For this purpose, we used the previously described score for the likelihood of SCM. Sequence-logos of the base following the stop codon position suggest that the presence of T after the stop codon significantly increases the protein synthesis termination efficiency (Fig 3C). On the contrary, G in the fourth position significantly increases the likelihood of SCM (Fig 3C). The identity of the base preceding the stop codon seems not to influence the likelihood of SCM (Fig 3D, see Method section for further details).

Overall, these analyses indicate that the downstream region after the premature stop codon influences the SCM rate. While G increases the SCM propensity, T has the opposite effect. It could be that the higher stability of the secondary structures in nucleotide regions of the mRNA with high GC content hampers the fidelity of the translation.

### Stop codon miscoding events relate to amino acid misincorporations

Next, we investigated the events occurring at the position of the stop codon in the cases where protein synthesis termination error was detected. We chose nine reporters for which SCM event was detected, and the premature stop codon was located within tryptic peptide detectable by mass spectrometry. The selected reporters were expressed in *E. coli* grown on nutritionally rich medium (LB) at 18°C; the expressed products were purified using C-terminal His-tag by Ni-NTA chromatography and analysed by GeLC-MS/MS. Wild-type mScarlet sample was processed similarly and used as a control.

Protein synthesis termination error may relate to the misincorporation of an amino acid at the position of the stop codon, skipping of the stop codon^37^, or a frameshift. Our data showed that misincorporation occurred for all nine selected reporters producing a mixture of several products, and the misincorporated amino acids were not a random selection (Fig 3D, Fig S6, and Table S1). TGA stop codon was reproducibly replaced by tryptophan and cysteine, and quantitative estimates suggest tryptophan as the main choice. Misincorporation of a few other amino acids at the TGA position was clearly a minor process (Table S1). On the other hand, TAG stop codon followed another pattern. It was repeatedly replaced by tyrosine, glutamine, and lysine, with all three amino acids having comparable misincorporation rates. TAA stop codon exhibited, in the only sample studied, replacement by tyrosine, glutamine, lysine, and alanine. No misincorporation events were detected in the control sample.

Overall, most of the detected amino acid misincorporations can be explained by the single nucleotide mismatch between the tRNA anticodon and the stop codon. Similarly, organisms that exhibit stop-to-sense reassignment of stop codons include TAG and TAA reassignment to Glutamine^38^ and TGA reassignment to Tryptophan^39^. Further, our results corroborate previous observations reported for *S. cerevisiae* where TAG and TAA were found to be replaced by glutamine, lysine, and tyrosine, and TGA - by tryptophan, cysteine, and arginine^40^. In mammals, tryptophan, cysteine, arginine, and serine can be incorporated at TGA stop codon position^37^.

We detected only a single mass spectrum which was matched to a peptide with a stop codon deletion (for the sample with TGA at position 190), suggesting that skipping the stop codon is a non-significant contributor to SCM events. No cases of extended deletion (stop codon together with 1-2 neighbouring amino acids) were detected. We also did not observe alternative-frame peptides, as expected, since we purified the proteins using a C-terminal tag that is in frame.

Altogether, our results indicate that SCM events are due to amino acid misincorporations, and the pattern of misincorporated amino acids depends on the stop codon identity.

### Low RNA polymerase accuracy at premature stop codons

We wondered whether the amino acid misincorporations identified by LC-MS/MS are due to transcription or translation errors. To answer this inquiry, we studied the mRNA sequence of 12 reporters, including the nine reporters previously studied by mass spectrometry and the mScarlet wild-type, expressed in wild-type *E. coli* grown at 18°C in LB media (Data S2). We confirmed with DNA sequencing the samples sequences (Data S3).

The most surprising aspect of the RNA-seq results is the higher probability of mismatches at the premature stop codon sites (Fig S7A). Even though some of these mismatches do not result in an amino acid exchange in the protein sequence due to the degeneration of the genetic code, i.e., several codons encode the same amino acid, it still looks like the errors at the premature stop codon sites are unusually high (Fig S7B). We thought that the reason for the RNA polymerase inaccuracy could be due to the nucleotide context around the premature stop codon. However, the probability of a mismatch in a given position is significantly higher when the nucleotide encodes for a premature stop codon than when it encodes for an amino acid or the canonical stop codon (Fig 4A). The same trend can be observed when considering only non-synonymous mismatches (Fig 4B). To further study the effect of the sequence context, we analysed the probability of a mismatch based on the adjacent nucleotides (Fig 4CD). This analysis revealed that the identity of the adjacent nucleotides does not influence the accuracy of the RNA polymerase. Interestingly, according to this analysis, the RNA polymerase is more prone to mismatch a stop codon when it is a premature one and it is in the ribosome’s open reading frame, which points to the coupling of translation and transcription as a putative reason for the higher inaccuracy of the RNA polymerase.

**Figure 4.**
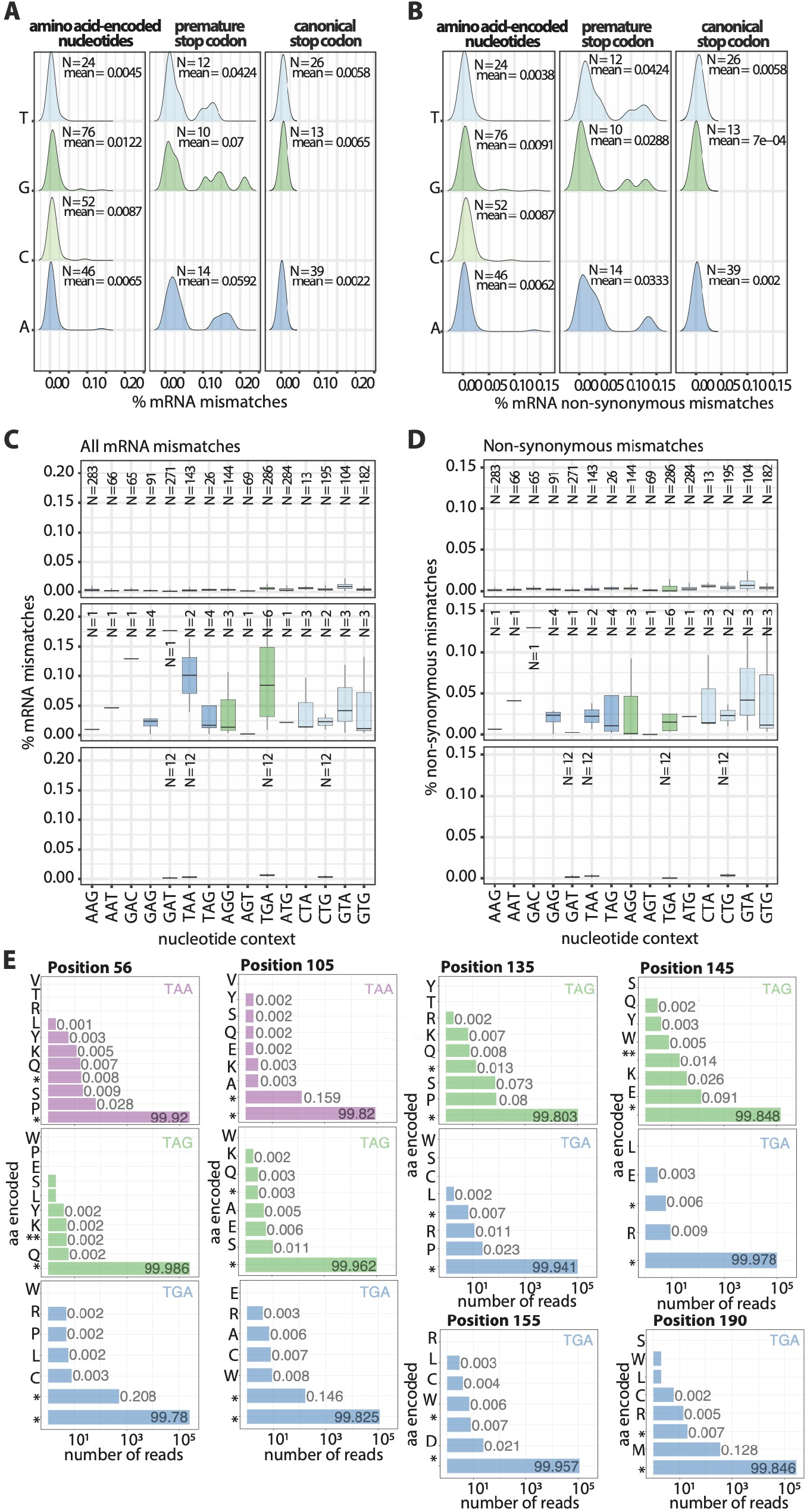
Low RNA polymerase accuracy at premature stop codons. **A)** The likelihood of the RNA polymerase mismatching is higher at a premature stop codon. **B)** The identity of the neighbouring nucleotides does not influence the accuracy of the RNA polymerase. Thus, the RNA polymerase is more prone to incorporate a mismatch at a stop codon only when it is in frame. **E)** RNA polymerase errors at premature stop codons result in a broad range of misincorporated amino acids.

We then moved to study whether the nucleotide misincorporations by the RNA polymerase errors match the protein sequences identified by MS and, in this way, clarify the source of the SCM: is it transcriptional or translational? We analysed the amino acids encoded by the mRNA sequences at the premature stop codon in each reporter (Fig 4E). As expected, the wild-type stop codon is found in most of the reporters’ mRNA sequences. Due to the degeneration of the genetic code, i.e., several codons code for the same amino acid or stop signal, RNA polymerase errors often result in a different stop codon, that is only different in one nucleotide (e.g., TAA to TAG) (Fig. 4BD). However, sometimes RNA polymerase errors result in an amino acid insertion at the premature stop codon site. The amino acid detected by mass spectrometry was often found already encoded in the mRNA sequences due to nucleotide misincorporations, although in a much smaller proportion and among many other amino acids. Overall, the RNA-seq and LC-MS/MS results suggest that mainly translation errors contribute to SCM events. Transcription errors contribute less to SCM events, yet they diversify the resulting protein sequence independent of the stop codon identity.

### Proteome-wide detection of stop codon miscoding events reveals the conditional expression of non-coding sequences in *E. coli*

To test whether and to what extent stop codon miscoding occurs in a natural context, we searched for evidence of SCM in wild-type *E. coli* proteome using untargeted mass spectrometry. Since growth temperature proved to be crucial for SCM with the reporter’s experiments, we grew clonal populations of *E. coli* at 37 and 18°C.

In total, we identified 17 peptides expressed from non-coding regions via SCM, mapping to 16 different proteins (Table S2). Of the 17 peptides, 13 covered the stop codon site and thus identified the inserted amino acid at the stop codon. For the ribosomal protein rpsG, we identified both Cys and Trp at the TGA site (Table S2 and Fig S8).

While we found that TAA can be miscoded by many amino acids, TGA is primarily miscoded by Trp (5 out of 10 cases, Fig 5A, Table S2). Since Trp is one of the rarest amino acids in the *E. coli* proteome, it seems overrepresented among the SCM peptides. The tendency to insert Trp is observed in both the fluorescent reporters and the proteome-wide study. In contrast, no preference was observed for the TAA stop codon.

**Figure 5.**
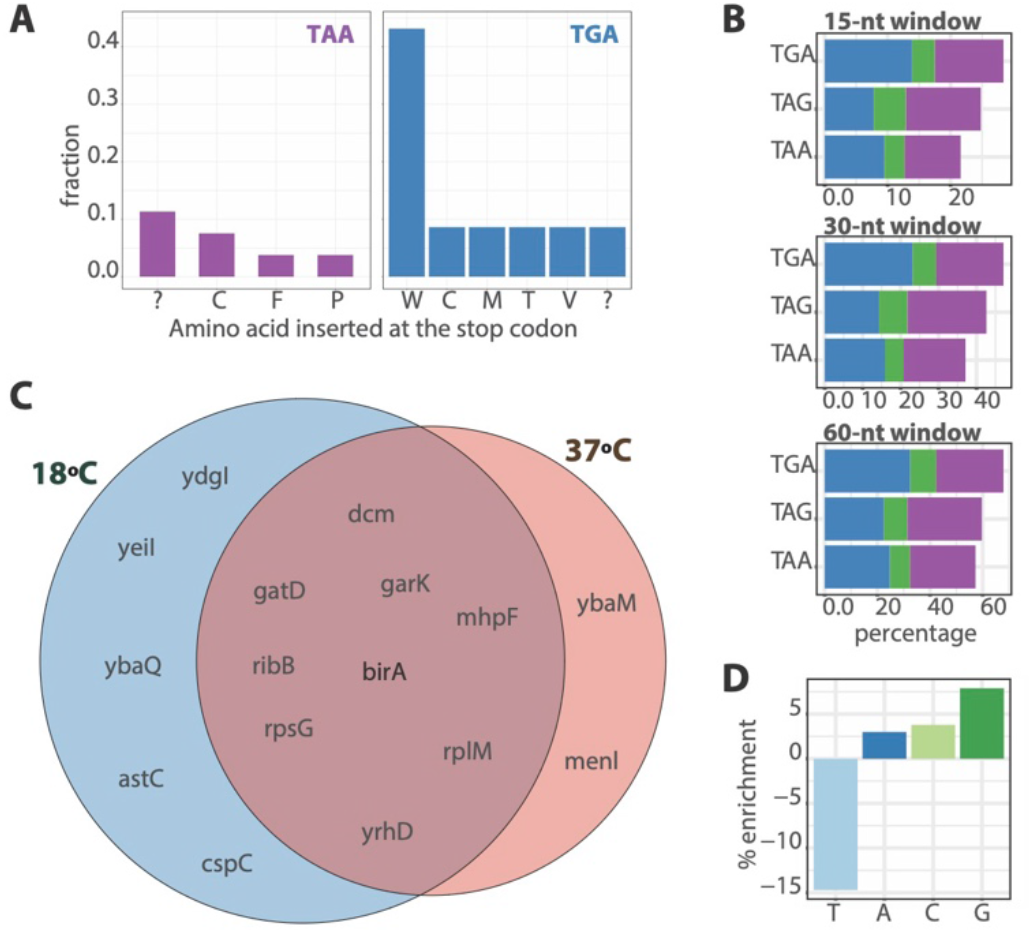
Proteome-wide mass spectrometry detects stop codon miscoding events in 32 peptides mapping to 28 proteins in *E. coli*. **A)** We detected SCM in 0.45% of TAA *E. coli* proteins and 1.35% of TGA *E. coli* proteins. While we found that TAA can be miscoded by many amino acids with similar frequencies, TGA is usually miscoded by Trp and Cys. **B)** The occurrence of an additional stop codon in a 15, 20, and 30-nt window downstream of the stop codon correlates with the protein synthesis accuracy of the stop codon in *E. coli* genome. TAG, as the least accurate stop codon, has the highest frequency of an additional stop codon in the 3’ regions of genes. **C)** We detected more cases of SCM in *E. coli* samples grown at l8°C than at 37°C. 9 genes were exclusively detected at 18°C and only three genes were detected at 37°C. **D)** The identity of the nucleotide downstream of the stop codon modulates the probability of SCM. While G decreases the protein synthesis termination accuracy, T increases the protein synthesis termination accuracy. Stop codon miscoding, SCM.

We observed a preference for SCM at TGA sites. Of the 16 genes where SCM was detected, 10 contain TGA, representing 0.86% of all TGA-containing *E. coli* proteins. However, only 6 genes contain TAA, representing 0.23% of TAA-containing *E. coli* proteins. We could not detect any evidence for SCM events at genes with TAG as the stop codon, probably due to its low representation (8%, Fig 5A). This is in line with the observation from the reporter library that TGA is the most error-prone of the three codons.

We then looked for additional stop codons in a 15-nt window downstream of the canonical *E. coli* stop codons. The probability of an additional stop codon correlates with the protein synthesis termination accuracy. For example, it is more likely to find an additional stop codon after TGA, the most error-prone stop codon, then after TGA, and then after TAA (Fig 5B). This trend is maintained if we extend the search to 20-nt and 30-nt windows (Fig 5B), suggesting differences in selection pressure to fix an additional stop codon depending on the accuracy of the stop codon.

In agreement with the fluorescence reporter’s study, we detected more cases of SCM in *E. coli* samples grown at l8°C than at 37°C (a peptide was considered present in a condition if it was identified in at least three out of six replicates. Five genes were exclusively detected at 18°C. In comparison, only two genes were detected at 37°C (Fig 5c, Table S2 and S8). Overall, this reinforces the conclusion drawn from the library experiments that lower temperature increases SCM events (Fig 5C).

Lastly, we wanted to analyse how the genome context modulates the likelihood of SCM proteome-wide. The previous experiments with the fluorescence reporters indicate that the identity of the base following the stop codon influences the likelihood of SCR (Fig. 3BC). While G seems to increase the likelihood of SCM, T seems to decrease the likelihood of SCM events. We then wondered whether the identified 16 genes with SCM follow the same trend. We calculated the enrichment of each nucleotide at the position immediately after the stop codon in the 16 identified genes compared with the rest of the genome. This study confirms that the identity of the nucleotide downstream of the stop codon modulates the likelihood of SCM. While G decreases the accuracy of the protein synthesis termination, T has the opposite effect (Fig 5D).

Overall, there is an agreement between the conclusions drawn from reporters and from the proteome-wide study. Furthermore, the proteome-wide mass spectrometry analysis revealed the expression of cryptic sequences –peptides from non-coding regions that can be generated only by SCM events– which are only expressed under certain environmental conditions.

## DISCUSSION

Here, we show that evolution frequently samples stop codon miscoding (SCM) events in *E. coli*. Furthermore, we report that internal factors, such as stop codon identity and genetic context, and external factors, such as growth temperature or nutrients, modulate the rate of SCM events (Fig. 2, Fig S1-S4). We also analysed the impact of the inaccuracy in protein synthesis termination on the proteome (Fig. 5, Table S2). Transcriptional errors on stop codons introduce a vast exploration of the mutational space since RNA polymerase miscodes premature stop codons often and towards different codons independently of the stop codon identity (Fig. 4E). Translational errors introduce a comparatively minor exploration of the mutational space depending on the stop codon identity (Fig. 3D, Table S1), yet they have significantly higher rate, being the main contributor to SCM events.

Despite the significant number of functional SCM reported^7,9,11–15 11^, most errors in protein synthesis termination are considered non-adaptative^17^. However, our findings show an error rate of up to 6% when bacteria is grown in normal conditions (Fig. 1E) (previous error rates recorded by fluorescence reporters for non-programmed SCM in *E. coli* grown under normal conditions were 2%^22,25^ and in *B. subtilis* 0.4%^23^). Even lower stop codon miscoding levels have often been linked to functionality^11^. For example, it has been reported how a 1% ribosomal readthrough level at a short conserved stop codon context is used in animals and fungi to generate peroxisomal isoforms of metabolic enzymes^14^. Further, our results indicate that the selection pressure to prevent SCM is not enough to reduce the usage of the most error-prone stop codon or increase release factors’ efficiency. Interestingly, wild-type *E. coli* K12 strains have an RF2 variant with reduced ability to terminate translation^41^. Instead, our data suggest selection pressure to fixate an additional stop codon downstream of the most error-prone stop codon (Fig. 5B).

Further, our data show that non-optimal growth temperatures and nutrient scarcity dramatically increase SCM events. As we observed in some of our reporters, SCM occurred at a rate up to 80% (Fig S2) (previous highest error rate for a non-programmed stop codon was 14% TGA readthrough when cells were grown in LB supplemented with lactose^22^). The effect of nutrient scarcity in SCM may be due to the lower carbon source concentration, which was previously linked to modulating the RF2 activity^22^. We propose that the effect of temperature on SCM could be explained due to the higher stability of the secondary structures of nucleotides at low temperatures. Strong downstream mRNA secondary structures hamper the unfolding of the mRNA, potentially influencing the accuracy of protein synthesis^42^. The same arguments can explain, on the one hand, why the downstream regions affect the protein synthesis accuracy while the upstream regions seem not to. On the other hand, it can also explain why G downstream the stop codon induces SCM since downstream stem-loops containing GC base pairs decrease the ribosomal accuracy^42^.

Based on these results, we propose avoiding TGA and optimizing the downstream region after the stop codon when designing a vector for protein expression in *E. coli* to minimize SCM events. The most common strategy to purify proteins involves *E. coli*, which is often grown at low temperatures, which may promote the production of non-desired protein forms^43^.

Overall, we hypothesize that bacteria could use errors in protein synthesis termination as a mechanism that allows the rapid diversification of its proteome to adapt to sudden environmental changes. We indeed detected cryptic sequences, among the *E. coli* proteome, that are only expressed by SCM under cold shock. Importantly, this analysis validated the proposed rules to predict endogenous SCM in vivo. Further studies are needed to elucidate the phenotypes and the potential functional role of these cryptic sequences.

We investigated the source of error behind these high SCM levels. We discerned that, although ribosomal errors are the main contributors to SCM, RNA polymerase errors contribute on a minor scale yet introduce a greater proteome diversification. Our study shows that RNA polymerase intriguingly misincorporated nucleotides in a non-random fashion. I.e., it mainly misincorporated at premature and in-frame stop codons. Future work will be needed to identify the molecular mechanisms that cause this bias in RNA polymerase error rate.

Our work highlights that both transcription and translation errors contribute to protein diversity. We showed that stop codon miscoding is more frequent than previously thought and it provides an evolutionary mechanism enabling cells to respond rapidly to the environment by increasing protein heterogeneity.

## ACKNOWLEDGEMENTS

We thank Andrej Shevchenko for his support and advice with mass spectrometry. We thank Lena Hersemann, Noreen Walker, and Andre Gohr from the Scientific Computing Facility of the MPI-CBG for helping with RNA-seq data and image analyses; Marc Bickle and Martin Stöter from the Technology Development Studio for guidance and assisting with the automatic microscope; Barbara Borgonovo, Eric Geertsma and Aliona Bogdanova from the Protein Biochemistry Facility of the MPI-CBG for helping with the western blot experiment; Julia Jarrells from the Cell Technology Facility of the MPI-CBG for the RNA extraction preparation and Sylke Winkler and Nicola Gscheidel from the Sequencing and Genotyping Facility of the MPI-CBG for the RNA-seq experiments.

## METHODS

### Gene Libraries

The reporter gene library was ordered from Twist Bioscience. The gene library was cloned in a pASK vector with a tetracycline-controlled promoter because it allows the tightly regulation of protein expression upon anhydrotetracycline titration^44^. We chose p15A origin of replication because it provides a low copy number of vectors in the cell^45^.

Below is the DNA and protein sequence of the wild-type mScarlet. In bold are highlighted the positions mutated to TAA, TAG, and TGA. In blue and red are highlighted the N-terminal Strep-tag and the C-terminal His-tag, respectively. Downstream of the His-tag we introduced two stop codons.

#### DNA sequence

ATGGCGAGCGCGTGGAGCCACCCGCAGTTCGAAAAAATGGTGAGCAAAGGC**GAA**GCGGTGATTAAA**GA A**TTTATGCGCTTT**AAA**GTGCACATGGAA**GGC**AGCATGAACGGC**CAT**GAATTTGAAATT**GAA**GGCGAAG GCGAA**GGC**CGCCCGTATGAA**GGC**ACGCAGACGGCG**AAA**CTGAAAGTGACG**AAA**GGCGGCCCGCTG**CCG** TTTAGCTGGGAT**ATT**CTGAGCCCGCAGTTTATGTATGGCAGCCGCGCGTTTACG**AAA**CATCCGGCGGA T**ATT**CCGGATTATTAT**AAA**CAGAGCTTTCCG**GAA**GGCTTTAAA**TGGGAA**CGCGTGATGAAC**TTT**GAAG ATGGCGGC**GCG**GTGACGGTGACG**CAG**GATACGAGCCTG**GAA**GATGGTACCCTG**ATT**TATAAAGTGAAA **CTG**CGCGGCACGAAC**TTT**CCGCCGGATGGC**CCG**GTGATGCAGAAA**AAA**ACGATGGGCTGG**GAA**GCGAG CACGGAA**CGC**CTGTATCCGGAA**GAT**GGCGTGCTGAAA**GGC**GATATTAAAATG**GCG**CTGCGCCTGAAA**G AT**GGCGGTCGCTAT**CTG**GCGGATTTTAAA**ACG**ACGTATAAAGCG**AAA**AAACCGGTGCAG**ATG**CCGGGC GCGTAT**AAC**GTGGATCGCAAA**CTG**GATATTACGAGC**CAT**AACGAAGATTAT**ACG**GTGGTGGAACAG**TA T**GAACGCAGCGAAGGCCGCCATAGCACGGGCGGCATGGATGAACTGTATAAACTCGAGCACCACCATC ACCATCACCATCACTGATAA

#### Protein sequence

MASAWSHPQFEKMVSKG**E**AVIK**E**FMRF**K**VHME**G**SMNG**H**EFEI**E**GEGE**G**RPYE**G**TQTA**K**LKVT**K**GGPL**P** FSWD**I**LSPQFMYGSRAFT**K**HPAD**I**PDYY**K**QSFP**E**GFK**WER**VMN**F**EDGG**A**VTVT**Q**DTSL**E**DGTL**I**YKVK **L**RGTN**F**PPDG**P**VMQK**K**TMGW**E**ASTE**R**LYPE**D**GVLK**G**DIKM**A**LRLK**D**GGRY**L**ADFK**T**TYKA**K**KPVQ**M**PG AY**N**VDRK**L**DITS**H**NEDY**T**VVEQ**Y**ERSEGRHSTGGMDELYKLEHHHHHHHH**

### *E. coli* strain and media

Plasmids encoding the reporters for SCM events were electrotransformed in the wild-type K-12 MG1655 *E. coli* strain. Transformants were grown in LB-agar plates and were inoculated into 384-well plates containing LB media supplemented with 15 µg/mL of chloramphenicol. To store the transformants at -80°C (glycerol stock), they were grown at 37°C to saturation, and glycerol was added until a 20% final concentration.

To investigate the temperature effect on SCM, *E. coli* cultures were grown into 384-well, without shaking, under saturated humidity conditions at 18, 25, 37, and 42°C to saturation. To address the nutrient depletion effect on SCM, LB, and M9 media were tested.

To titrate the expression of the Gly95-TGA reporter, 0, 25, 50, 100, 200, 400, and 800 µg/L of anhydrotetracycline was added to the media. For the study of the library, to induce the expression 400 µg/L of anhydrotetracycline was added to the media. To avoid the photodegradation of the anhydrotetracycline, cells were grown under light protection.

#### M9 standard media

M9 media supplemented with 0.4% glycerol, 0.2% casamino acids, 1 mM thiamine hydrochloride, 2 mM MgSO_4_ and 0.1 mM CaCl_2_.

#### M9 supplemented with higher carbon source concentration

M9 media supplemented with 1.6% glycerol, 0.2% casamino acids, 1 mM thiamine hydrochloride, 2 mM MgSO_4_ and 0.1 mM CaCl_2_.

#### M9 supplemented with higher casamino acid concentration

M9 media supplemented with 0.4% glycerol, 0.4% casamino acids, 1 mM thiamine hydrochloride, 2 mM MgSO_4_ and 0.1 mM CaCl_2_.

### Microscopy Screenings and Sample Preparation

A liquid handling robot (Beckman Coulter Biomek FXp with Thermo Cytomat 6002) was used to perform plate-to-plate transfers of cells. The cells were inoculated from glycerol stock into 384-well plates (Eppendorf microplate 384/V #0030621301) containing 100 µL of LB-media supplemented with 15 µg/mL of Chloramphenicol. Preculture plates were then grown until cells reached saturation (∼24h) at 37°C. Then, 2µL of culture from the saturated cultures were used to inoculate the 384-well plates (Eppendorf microplate 384/V #0030621301) containing 100 µL of the appropriate media (LB or M9) supplemented with 15 µg/mL of Chloramphenicol and 400 µg/L of anhydrotetracycline. The cells were grown at different temperatures (18, 25, 37, and 42°C) until they reached saturation (24-48h) under light protection. Then, cells from the saturated cultures were transferred into Greiner 384-well glass-bottom optical imaging plates (#781092) previously coated with poly-L-lysine containing 50 µL of PBS. To coat the Greiner 384-well glass-bottom optical imaging plates, 50 µL of 0.01% (w/v) of poly-L-lysine (SIGMA, #P4832) was added and incubated for at least 1 h. Then, the plates were washed with water and were dried overnight.

All confocal imaging was performed on an automated spinning disc confocal microscope (CellVoyager CV7000, Yokogawa), using a 60x 1.2NA objective. For excitation, a 561nm laser was used, and the fluorescence was detected through a 600/37 band pass emission filter. We recorded 2560x2560 16-bit images at binning 2 for brightfield and for mScarlet fluorescent protein in epifluorescence mode on sCMOS cameras. A laser-based hardware autofocus was used to acquire 5 x to 9 images per well.

### Fluorescence data analysis

Image analysis was done with Fiji^46^ and downstream data analysis and visualization was performed using R v4.1.2.

The cells were identified and segmented, and their fluorescent signal (mean and standard deviation, mode, minimum, and maximum) as well as additional cell properties (area, x- and y-coordinates) were determined in Fiji using custom macros^46^ (fluorescence data per cells is provided as Data S1).

### Western Blot

Single-colony *E. coli* LB-cultures were grown at 37°C until an absorbance of 0.6 at 600nm. Then, they were grown at 18°C for 2h. Protein expression was induced by addition of anhydrotetracycline to a final concentration of 400 µg/L, and cells were grown overnight at 18 °C. Aliquots of around 1mg of total proteins (A_600nm_ = 1≈ 0.3 mg/mL) were centrifuged. Cell pellets were resuspended in 400µL of ice-cold disruption buffer (PBS containing 10% Glycerol, 1mM MgSO_4_, Benzonase 0,05U/m, Roche complete cocktail EDTA free 1 tablet/10mL) and 300 mg glass beads (0,1 mm Scientific industry SI-BG01) were added. Cells were disrupted in FastPrep-24™ (MP Biomedicals) at low temperatures and centrifuged.

Supernatants were processed by SDS–polyacrylamide gel electrophoresis (4–20 % Tris-Glycine-Gel, Anamed #TG 42015).

All gels contain the lysate of cells carrying the mScarlet (PC), an empty vector (NC) and a protein marker. The mScarlet and marker bands were visualized with a Typhoon 9500 at 532 nm directly from the gel. The rest of the proteins were transferred to nitrocellulose membranes (Whatman BA85) performing semi-dry blotting (Transfer buffer: 20 mM Tris-Base, 160 mM Glycine, 0,1% SDS, 20 % Methanol). From the membranes, total protein amounts were assayed using Fast Green FCF (Sigma, #F7252) staining solution^47^ and imaged using LICOR Odyssey 700 nm. For immunodetection the membranes were blocked with Blocking buffer (5% milk, 0,1% Tween20 in PBS) followed by incubation with 1:5000 Qiagen mouse anti-penta His-tag (# 34660) and 1:10.000 Licor IRDYE 800 goat anti mouse (#926-32350). Signals were detected with LICOR Odyssey and analyzed with Fiji^46^.

The mScarlet sample (PC) shows in-gel fluorescence in several bands with a molecular weight around 26 kDa, probably due to degradation. The area selected to analyze the His-tag expression was the same where the PC presented fluorescence (Fig S5).

### Analysis of the influence of the genetic context

#### Stop codon miscoding likelihood score

Those reporters that displayed a median fluorescence below a threshold (defined as the mean plus two standard deviations of the fluorescence signal of the NC) got a score of 0. The reporter with the highest fluorescence signal was assigned the highest score with a value of 1.0. All other reporters were ranked between 0 and 1.0 according to their median fluorescence values. We repeated this ranking strategy for all the stop codons at all the tested conditions. Then, we calculate the average score among conditions and the three stop codons (24 values per position) to have a unique score per position and, therefore, per genome context.

#### Correlation between nucleotide content and stop codon miscoding likelihood score

The Stop codon miscoding likelihood scores were binned in 8 equal-bin-width. The number of samples per bin are: N= 12 (bin = 0), N=12 (0<bin<0.125), N=9 (0.125<bin<0.250), N=2 (0.250<bin<0.375), N=1 (0.375<bin<0.5), N=0 (0.5<bin<0.625), N=1 (0.625<bin<0.750).

#### Correlation between the nucleotide identity adjacent to the stop codon and stop codon miscoding likelihood score

The stop codon miscoding likelihood scores were binned into three categories: i) accurate protein synthesis termination (score = 0, N = 12). ii) medium tendency to SCM (0 < score < mean of all scores, N = 16); iii) high tendency to SCM (score > mean of all scores, N = 9).

### Protein purification

Wild-type mScarlet for the stability assay, for the calibration curve, and the samples to study by mass spectrometry were expressed in *E. coli* and purified with His-tag affinity chromatography. Briefly, genes encoding these proteins with C-terminal 8x His-tag were cloned into a pASK vector under tetracycline-controlled promoter. Vectors were transformed into K-12 MG1655 *E. coli* cells. From this point, two procedures were used to grow the cells and induce the expression of the proteins: i) To purify the samples for mass spectrometry analyses, cells were grown on LB-medium supplemented with 15 µg/mL of chloramphenicol at 37°C until absorbance of 0.2 at 600nm, then at 18°C until reaching absorbance of 0.6 at 600nm. Protein expression was induced by anhydrotetracycline (final concentration of 400 µg/L) and cells were grown overnight at 18°C (∼12h). ii) To purify the wild-type mScarlet for the stability assay and for the calibration curve, cells were grown on LB-medium supplemented with 15 µg/mL of chloramphenicol at 37°C until absorbance of 0.6 at 600nm. Then, protein expression was induced by the addition of anhydrotetracycline (final concentration of 400 µg/L) and cells were grown for 4h at 37 °C.

Cells were harvested by centrifugation and resuspended in lysis buffer comprising 20 mM sodium phosphate, 500 mM NaCl and 20 mM imidazole, pH 7.4 supplemented with Roche complete cocktail EDTA free 1 tablet/10mL and benzonase 0,05U/m. Cells were lysed using LM20 microfluidizer (Microfuidics). The lysates were clarified by centrifugation for 1h at 4ºC at 12000 rpm and loaded on a His GraviTrap™ column (GE Healthcare). After washing with 10 mL of washing buffer (40 mM sodium phosphate, 500 mM NaCl and 20 mM imidazole, pH 7.4), proteins were eluted with sodium 20 mM sodium phosphate, 500 mM NaCl, and 500 mM imidazole, pH 7.4^48^.

The purity of all the samples was assessed by SDS–polyacrylamide gel electrophoresis (Life Technologies GmbH) and protein concentration was determined by absorbance at 280 nm (using the extinction coefficient of the wild-type mScarlet for all the samples, 39880 M^-1^cm^-1^). For the mass spectrometry analyses 50μg of each sample was separated by SDS-PAGE and mScarlet was purified for the thermostability assay and for the calibration curve was dialyzed against PBS for subsequent analyses.

### Calibration curve for fluorescence measurements

Triplicates of purified mScarlet were fluorescence imaged at 0, 3, 4, 7, 10, 15, 23, 35, 52, 78, 117, 175, 262, 393, 590, 885, 1327, 1991 and 2986 nM in PBS. Saturation of the fluorescence signal above 70000 AU defines the upper limit of the dynamic range (Figure S1). A linear relation between mScarlet concentration and fluorescence arbitrary units was observed between 40 and 800 nM.

### mScarlet thermostability assay

Aliquots of purified mScarlet at 60nM in PBS were incubated at a range of temperatures (30.5 to 98.2ºC) for 30 min. Samples were then fluorescently imaged. The procedure was performed in triplicates.

Unfolded mScarlet will not be functional and, therefore, will not show fluorescence. On the contrary mScarlet that remains folded, will show fluorescence. The thermostability curve (Fig S3C) indicated that mScarlet remained functional, i.e., fluorescent until 70°C.

### Mass Spectrometry analysis of reporters

Gel regions corresponding to the molecular weight of the reporters were excised and analysed by LC-MS/MS. Briefly, samples were in-gel digested with trypsin (sequencing grade, Promega, Mannheim), the resulting peptides extracted by two changes of 5% formic acid (FA) and acetonitrile, and dried down in a vacuum centrifuge. Peptide pellets were dissolved in 100μL of 5% FA and 5μL aliquot of peptide mixture was taken for MS analysis.

LC-MS/MS analysis was performed on a nanoUPLC Vanquish system interfaced on-line to an Orbitrap HF hybrid mass spectrometer (both Thermo Fischer Scientific, Bremen). The nano-LC system was equipped with Acclam PepMap™ 100 75 µm x 2cm trapping column and 50cm μPAC analytical column (Thermo Fischer Scientific, Bremen). Peptides were separated using 75min linear gradient, solvent A - 0.1% aqueous FA, solvent B - 0.1% FA in acetonitrile. Samples were first analyzed using data-dependent acquisition (DDA) and then by targeted acquisition with inclusion list guided by the results of RNA-seq analysis. DDA analysis was performed using Top20 method, precursor m/z range was 350-1600; mass resolution (FWHM) – 120 000 and 15 000 for MS and MS/MS spectra respectively; dynamic exclusion time was set on 15s. The lock mass function was set to recalibrate MS1 scans using the background ion (Si(CH3)2O)6 at m/z 445.1200. Targeted analysis was performed in profile mode, a full mass spectrum at the mass resolution of 240 000 (AGC target 3×10^6^, 150 ms maximum injection time, m/z 350–1700) was followed by PRM scans at the mass resolution of 120 000 (AGC target 1×10^5^, 200 ms maximum injection time, isolation window 3 Th) triggered by a scheduled inclusion list. To avoid carryover, 3-5 blank runs were performed after each sample analysis, the last blank was recorded and also searched against a customized database.

Spectra were matched by MASCOT software (v. 2.2.04, Matrix Science, UK) against a customized database comprising *E. coli* protein sequences extracted from UniProt database (version October 2022), and a set of modified mScarlet sequences. mScarlet sequences included three-frame translated nucleotide sequences with stop codons insertions, sequences with deletion of 1-2 amino acids surrounding the position of the stop codon that were denoted as “X” (equivalent to any amino acid). Database search was performed with 5ppm and 0.025Da mass tolerance for precursor and fragment ions respectively; enzyme specificity – trypsin; one miscleavage allowed; variable modifications – methionine oxidation, N/Q deamidation, cysteine sulfonic acid, cysteine propionamid, peptide N-terminal acetylation. The results were then evaluated by Scaffold software (v.4.11.1, Proteome Software, Portland) and also manually inspected. Identification of modified peptide was accepted if passed 95% peptide probability threshold and if the matched fragmentation spectra (minimal number of PMS: 2) comprised fragment ions unequivocally confirming the misincorporated amino acid (Fig. S6). Ratio of peptide forms comprising misincorporated amino acids was estimated based on extracted ions chromatograms (XIC) for each form produced by XCalibur Qual Browser, (Thermo Fischer Scientific) and normalized to the sum intensity of all forms of the peptide. Peptides with misincorporated Lys were excluded from the calculation.

### RNA-seq

Single-colony *E. coli* LB-cultures were grown at 37°C until an absorbance of 0.4 at 600nm. Then, they were grown at 18°C until an absorbance of 0.6 (∼2h), protein expression was induced by addition of anhydrotetracycline to a final concentration of 400 µg/L, and cells were grown overnight at 18 °C. Cells were diluted to an absorbance of 1.0 at 600nm and 100μL of of the cell suspension was mixed with with 200μL of RNAprotect bacteria reagent (Qiagen #76506). Cells were harvested by centrifugation and pellets were frozen. Then, total RNA of the samples was extracted according to manufacturing specifications (RNeasy Protect Bacteria Mini Kit Qiagen kit).

300 ng of total RNA per sample was mixed with 50 ng of random hexamers and dNTPs and hybridized at 65°C for 5 minutes. cDNA was reversely transcribed making use of the Thermo Fisher Superscript IV transcriptase according to the manufacturer’s instructions. Then, mScarlet was specifically amplified from the resulting cDNA with forward primer 5’ AGTTATTTTACCACTCCCTATCAGT 3’ and reverse primer 5’ AGTAGCGGTAAACGGCAGAC 3’ resulting in an amplified PCR fragment of 948 bps. The NEB Q5® High-Fidelity DNA polymerase was used according to the manufacturer’s instructions. PCR conditions were as such, an initial denaturation step for 30 sec at 98°C, 30 cycles of 10 sec denaturation at 98°C followed by annealing at 67°C for 30 sec and extension at 72°C for 30 sec. A final extension step was done for 2 min at 72°C. Primer and dNTPs have been removed with 1x volume AMPure bead purification. Amplified mScarlet fragments were finally quantified with the Thermo Fisher Qubit high sensitivity DNA quantification system. 1 ng of the amplified mScarlet fragments were analyzed on the Agilent Fragment Analyzer system making use of the NGS high sensitivity kit.

Pacbio SMRTbell® libraries have been generated following the PacBio® Barcoded overhand adapters for multiplexing amplicons (Express template kit 2.0). For each multiplexed library, eight samples have been pooled equimolarly. In brief, 50 ng of each amplified mScarlet fragments was damage repaired, followed by end repair and A-tailing according to the instructions. Pacbio barcoded overhang adapters (BAK8A and BAK8B) were ligated to the PCR fragments and equimolarly pooled prior to two final AMPure bead purification steps (1x volume). The final quality control of the resulting library was done on the Agilent Fragment Analyzer with the large fragment kit. We generated in total, two different 8plex PacBio HiFi libraries. The v4 PacBio sequencing primer in combination with the SEQUEL II binding kit 2.1 was used to sequence both eightplex PacBio SMRTbell® libraries. 80 pM and 120 pM of each library was loaded by diffusion loading, pre-extension time was 0.3 hours and run time 10 hours on the SEQUEL II making use of the SEQUEL II sequencing 2.0 chemistry. Circular consensus reads have been called with the PacBio SMRT link ccs calling tool and they were demultiplexed with lima, the PacBio demultiplexer and primer removal tool (https://lima.how/).

The long PacBio RNA-seq reads were mapped to the reference with BWA v0.7.17-r1198. BAM files representing the mapped reads were further processed with Samtools mpileup v1.15.1 using htslib 1.15.1 to generate a textual description of the mapped reads, including information about at which positions and reads mutations, insertions, deletions, and indels were found (all BAM files are provided as Data S2). The pileups were further processed with R to extract information on frequencies of mutated trimers along the mScarlet gene. For extracting information on synonymous and non-synonymous mutations PySam v0.16.0 with Python 3.7.12 was used with the BAM/SAM output from BWA. All plots were generated with base plotting of R v4.1.2.

### Untargeted Detection of Stop-Codon Miscoding in *E. coli*

#### Cell culturing and sample preparation

*E. coli* (strain K-12 MG1655) was grown overnight at 37°C on LB medium, then split in two parts, one incubated at 37°C, the other at 18°C until reaching OD600 of ca. 0.6. The experiment was performed in three biological replicates, samples were prepared and measured in a block-randomised fashion^49^ with two technical repeats. Cells were pelleted by centrifugation at 4,500g for 10 minutes, washed with PBS, and resuspended in 1ml of lysis buffer consisting of 8M Urea, 0.1M ammonium bicarbonate, 0.1M NaCl, and 1x Roche cOmplete™ Protease Inhibitor Cocktail (Roche Diagnostics Deutschland GmbH, Germany). Then 2 micro spatula spoons of 0.5 mm stainless steel beads (Next Advance Inc., USA) were added, cells lysed in a TissueLyser II (QIAGEN GmbH, Germany) for 2x 5min at 30Hz at 4ºC and the debris removed by centrifugation for 10min at 13000 g. Protein concentration in the supernatant was measured by Pierce BCA Protein Assay (Thermo Scientific, USA) and aliquots of 100 µg of proteins were taken for LC-MS/MS analysis. After reduction and alkylation, proteins were precipitated with isopropanol^50^ and digested overnight at 37°C with 1:50 enzyme : protein ratio by Trypsin/Lys-C Mix (Promega GmbH, Germany). Resulting peptides were desalted on MicroSpin column (The Nest Group, Inc., USA) and dried down in a vacuum concentrator. Prior mass spectrometric analysis, the samples were reconstituted in 0.2% aqueous formic acid, peptide concentration was determined by measuring absorption at 280 nm and 260nm using Nanodrop 1000 ND-1000 spectrophotometer (Thermo Fisher Scientific Inc, USA) and the Warburg-Christian method^51^ and adjusted to final concentration of 0.12 µg/µL; 5 µL were then taken for analyses.

#### LC-MS/MS analysis

LC-MS/MS analysis was carried out on the mass spectrometric equipment described in **Mass Spectrometry analysis of reporters** section. Peptides were separated using 120min 2-sloped gradient, solvent A - 0.1% aqueos FA, solvent B - 0.1% FA in acetonitrile: 80min 0 to 17.5 % ACN, 40min 17.5 % to 35% ACN at a flow rate of 0.5 µl/min. The gradient was followed by a 7min wash with 95% ACN. To avoid carryover, 2 blank runs were performed after each sample analysis. Further settings: spray voltage - 2.5kV, capillary temperature - 280°C, S-lens RF value - 50. Spectra were acquired by Data Independent Acquisition (DIA) in a staggered fashion^52,53^; full-scan mass spectrum with a mass range of 395-971 m/z and resolution 60,000 (AGC 3e6, 40 ms maximum injection time, fixed first mass 100) was followed by 32 MS2 scans in centroid mode at mass resolution 30,000 with isolation window 18 m/z covering mass range 400-966 m/z (55 ms maximum injection time, AGC 1e6, normalized collision energy 24%).

#### Database search and validation of candidate peptides

Acquired spectra were matched against customized database using DIA-NN software suit v1.8^54^. Customized database including sequences resulting from potential stop codon miscoding (SCM) events was created based on *E. coli* K12 MG1655 reference genome and the corresponding genome annotation in NCBI database (accession GCF_000005845.2, version from 18/05/21). For each gene, the ‘downstream sequence’ was determined by finding the next in-frame ORF after that stop codon in the genome (using the reverse complement for genes on the negative strand). The minimum length of the downstream sequence was 60 nucleotides, and the maximum was either 300 nucleotides or the distance to the next in-frame ORF. Genes were then *in silico* translated 20 times from canonical start to the end of the downstream sequence or until the second in-frame stop codon, each time translating the canonical (the first) stop codon with a different amino acid, and added to the fasta file. For each gene, the sequence list thus contained 20 sequences, each encompassing the canonical and genomic downstream sequence, with one of the 20 amino acids in place of the canonical Stop. This file as well as a file containing common contaminant sequences^55^ were used as input for DIA-NN to create the spectral library. Database preparation and data analysis was done in Python 3 and Jupyter Notebook, using pandas^56^, numpy^57^, and Biopython^58^.

Staggered DIA raw files were converted to .mzML format and demultiplexed using Proteowizard MSConvert v3.0.2^59^, and analysed with DIA-NN v1.8. Predicted spectral library was generated from sequences in customized database under the following settings: enzyme - Trypsin/P, one missed cleavage allowed; C(carbamidomethyl) as fixed modification; M(oxidation) and N-terminal M-excision as variable modifications, allowing up to one variable modification per precursor. Precursor and protein group matrices were filtered at 1% FDR. Other settings were: --double-search --smart-profiling --no-ifs-removal --no-quant-files -- report-lib-info --il-eq

The precursor-by-sample-matrix output generated by DIA-NN was processed further to identify SCM events. First, all precursors mapped to a contaminant protein or any sequence in Swissprot database (version from 22/12/21) were discarded. Remaining precursors mapped to the downstream sequence of an *E. coli* gene either fully (complete sequence is past the Stop codon) or partially (overlapping the Stop codon) and were treated as candidate SCM precursors. Singly charged precursors containing lysine or arginine were removed as well as precursors not extending for at least 2 amino acids over the stop codon. Further, only candidate SCM precursors reported in 3 out of 6 replicates of at least one temperature condition were retained. Fragmentation spectra of candidate SCM precursors were then manually validated in Skyline ‘daily’, version 22.2.1.351^60^. For this, the spectral library generated by DIA-NN based on observed fragment intensities (with ‘full-profiling’ library generation setting) and the peak boundaries from the DIA-NN main output table were imported into Skyline, as well as sequences of SCM precursors remaining after the described filtering. Candidates were judged by comparison of the empirical library spectra generated by DIA-NN against Prosit-predicted spectra^61^ as well as assessing chromatographic coelution of fragments. Peptide identification was accepted if i) peptide sequence matched at least three fragments (y3 and higher for tryptic peptides, b3 and higher or a combination of b/y-ions for other); ii) the fragments co-elute; iii) two fragments predicted by Prosit to be most intense (excluding y1, y2, b1, b2) had to be among matched. Peptides identified under these conditions were further considered for analyses of differential abundance, sequence context and Stop codon usage. To identify statistically significant differences in abundance between the 37°C and 18°C conditions, Student’s two-sided t-test with Benjamini-Hochberg adjustment^62^ was carried out on the log2 peptide intensities of the two groups. In cases where multiple precursors represented the same peptide sequence, the respective intensities were summed.

**Statistical analysis and data visualization** was performed by R v4.1.2. For all box plot representations thick black line indicates median, box indicates 25^th^ and 75^th^ percentiles, whiskers indicate 1.5 times the inter-quartile range. Figure 1A and D were created with Biorender (biorender.com) and Adobe Illustrator.

## DATA AVAILABILITY

The mass spectrometry data have been deposited in MassIVE repository under accession number MSV000091065 (Login: MSV000091065_reviewer and password: k64sa21). The raw fluorescent data obtained from microscopy images are provided as Data S1. RNA-seq data are provided as Data S2 (BAM files) and their DNA-seq chromatograms as Data S3 (bl1 files).

## Notes

### Competing Interest Statement

The authors have declared no competing interest.

