## Supplementary material for "Environment Modulates Protein Heterogeneity Through Transcriptional and Translational Stop Codon Miscoding": SUPLEMENTARY_MATERIAL

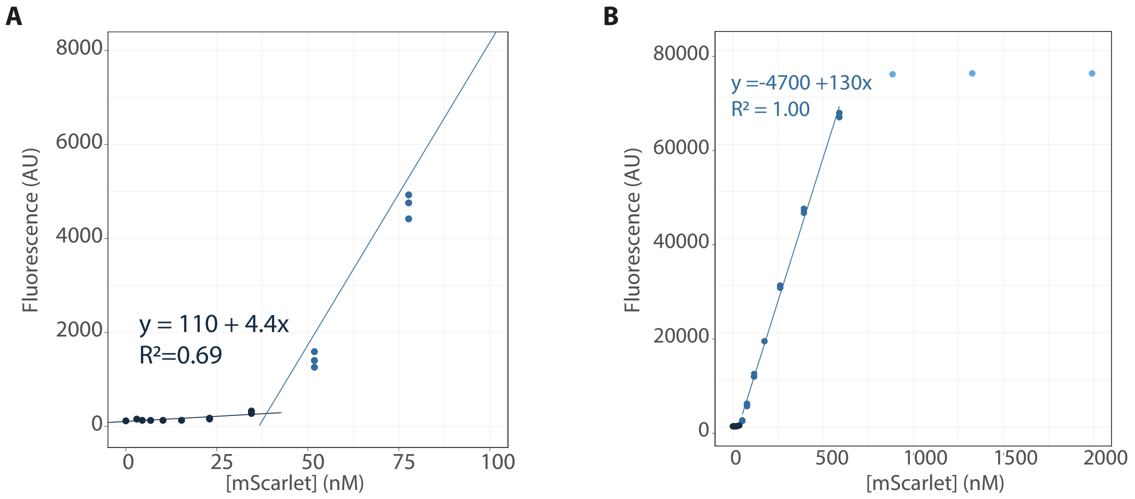


**Fig S1. Fluorescence measurements are within the linear dynamic range of the microscope (270-7000 AU). A)** Range of the calibration curve of mScarlet from 0-100 mM. The minimum concentration of mScarlet that we can determine with fluorescence measurements is 38 nM. **B)** Full range of the calibration curve of the mScarlet, from 0 -2000 mM. Saturation of the fluorescence signal above 70000 AU defines the upper limit of the dynamic range.

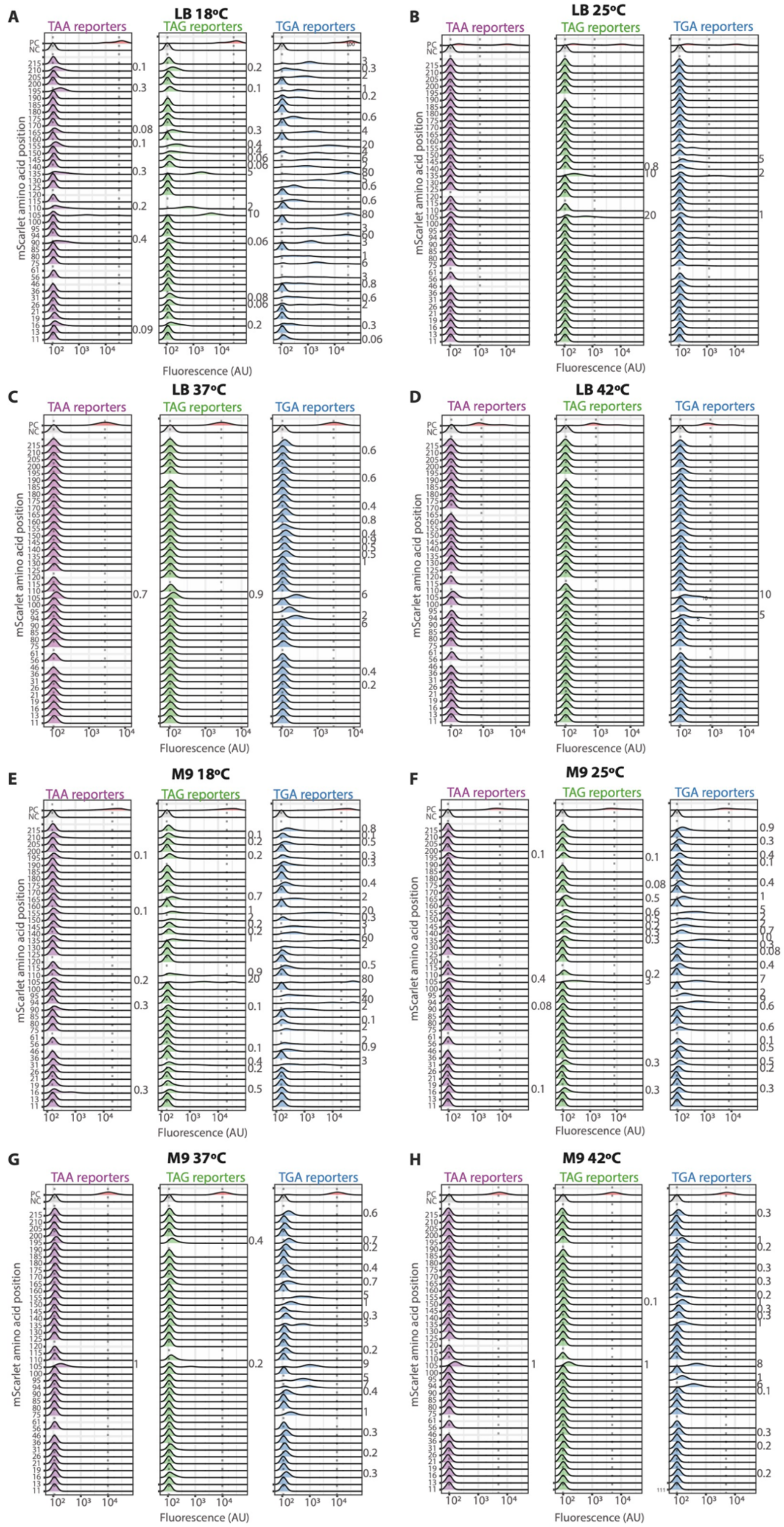


**Figure S2. Visualization and quantification of stop codon miscoding events in *E.coli* using fluorescence reporters.** Fluorescence intensity distributions displayed by the *E.coli* cells, transformed with each library's reporters and grown at: **A)** 18°C in rich media, **B)** 25°C in rich media, **C)** 37°C in rich media, **D)** 42°C in rich media, **E)** 18°C in minimum media, **F)** 25°C in minimum media, **G)** 37°C in minimum media and **H)** 42°C in minimum media. The propensity of SCM, calculated as the percent of the median fluorescence compared with the positive control, is shown for the distributions with a median fluorescence higher than the negative control. Non-optimal growth temperature and nutrient scarcity increase protein synthesis inaccuracy. Rich media: LB and minimum media: M9 media supplemented with 0.4% glycerol, 0.2% casa amino acids, 1mM thiamine hydrochloride, 2 mM $MgSO_4$ and 0.1 mM $CaCl_2$. The fluorescent data per cells are provided in Data S1.

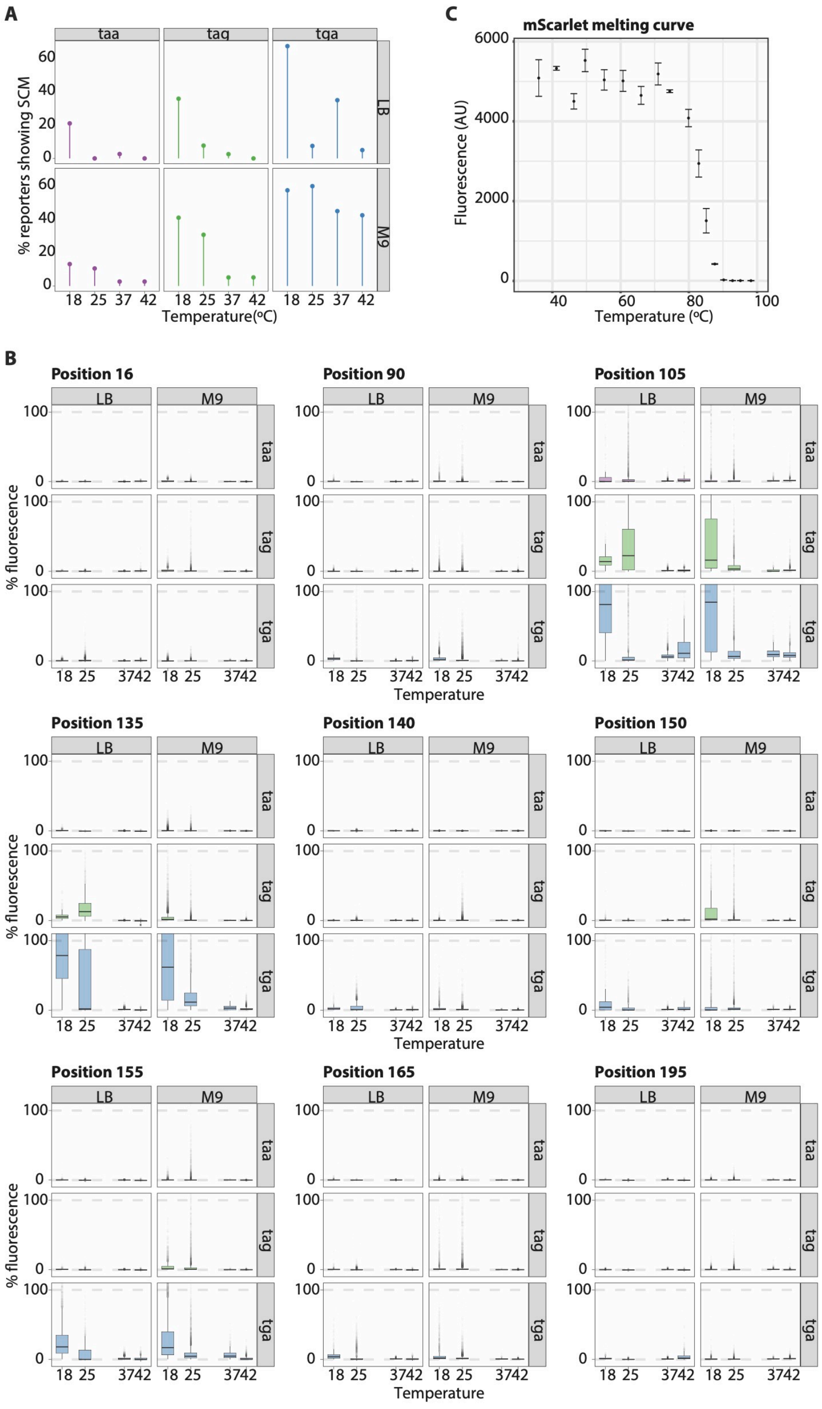


**Figure S3. Non-optimal growth temperatures and nutrient scarcity promote stop codon miscoding (SCM). A)** At low temperature more reporters display SCM, that was defined as having a median fluorescence higher than the threshold defined as the median fluorescence plus two standard deviation of the NC. More TGA reporters display SCM than TAG and TAA reporters. **B)** Fluorescence distributions summarized as box plots displayed by the *E. coli* cells of 9 selected reporters grown at different growth conditions. At these most error-pone positions: i) cells grown in minimum media (M9) dispalyed more errors than in rich media (LB). ii) TAA was the most accurate codon terminating the protein synthesis and TGA the least. iii) Cells grown at low temperature and cells grown in minimum media displayed more SCM events. **C) mScarlet thermostability assay.** mScarlet remained functional, i.e., fluorescent, until 70°C (mean and standard deviation of three replicates are shown).

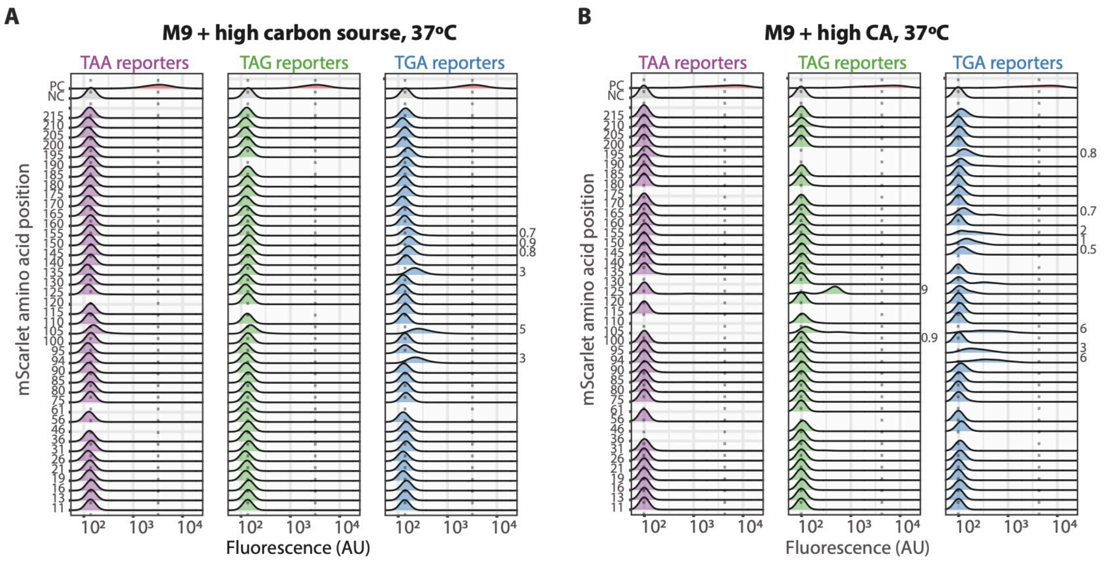


**Figure S4. Visualization and quantification of stop codon miscoding events in *E. coli* grown in different media.** Fluorescence distributions displayed by *E.coli* cells transformed with each library's reporters and grown at 37°C in minimun media supplemented with **A)** high carbon source concentration (M9 media with 1.6% glycerol, 0.2% casamino acids, 1mM thiamine hydrochloride, 2 mM MgSO 4 and 0.1 mM CaCl 2) and **B)** high casamino acid concentration (M9 media with 0.4% glycerol, 0.4% casamino acids, 1mM thiamine hydrochloride, 2 mM MgSO 4 and 0.1 mM CaCl 2).

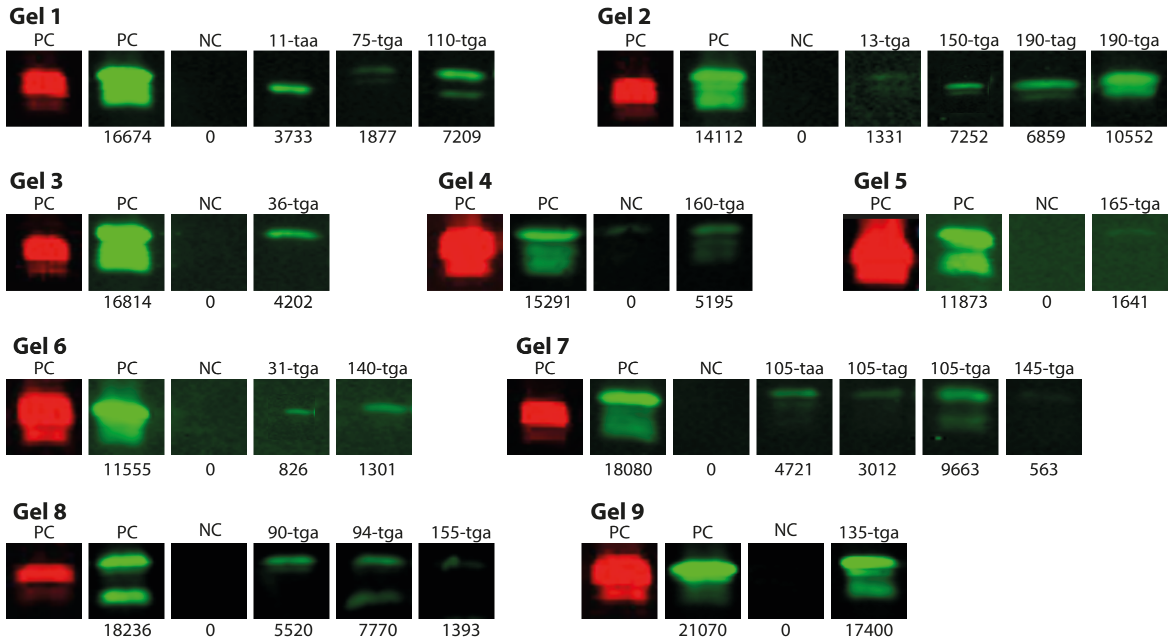


**Figure S5. Detection of stop codon miscoding quantifying the His-tag expression with His-tag antibodies.** In-gel fluorescence signals of the PC (cells expressing the mScarlet) are shown in red. The His-tag signal of the positive control (PC), and negative control, (NC, cells carrying an empty vector), and the cells expressing the reporters are shown in green. The numbers below the images indicate the quantification of the His-tag expression as explained in Method section. Cells were grown in rich media at 18°C (see Methods).

**Ala-105-TAA**

(R)VMNFEDGG**A**VTVTQDTSLEDGTLIYK(V)

sissi_rep_3_5_100ul_01.11519.11519.2.dta

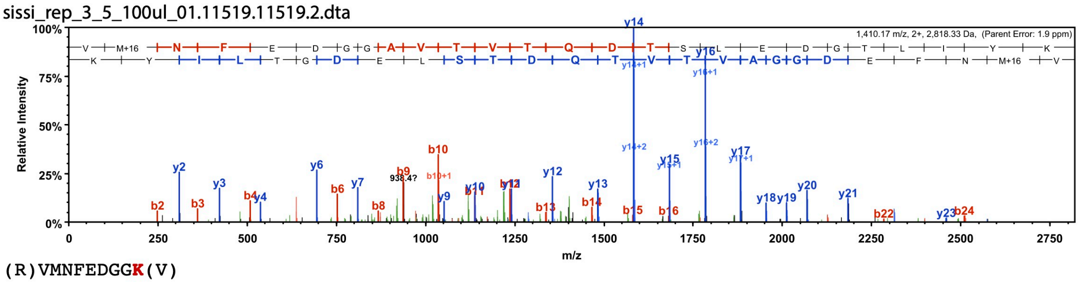


(R)VMNFEDGG**K**(V)

sissi_5_rep3_100ul.02204.02204.2.dta

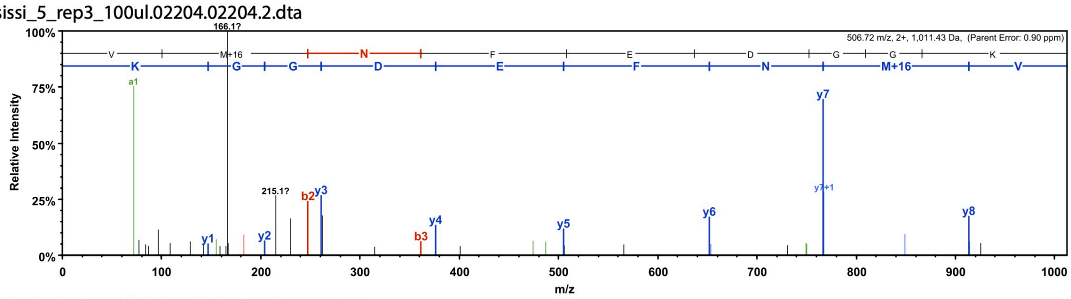


(R)VMNFEDGG**Q**VTVTQDTSLEDGTLIYK(V)

sissi_rep_3_5_100ul_01.10879.10879.2.dta

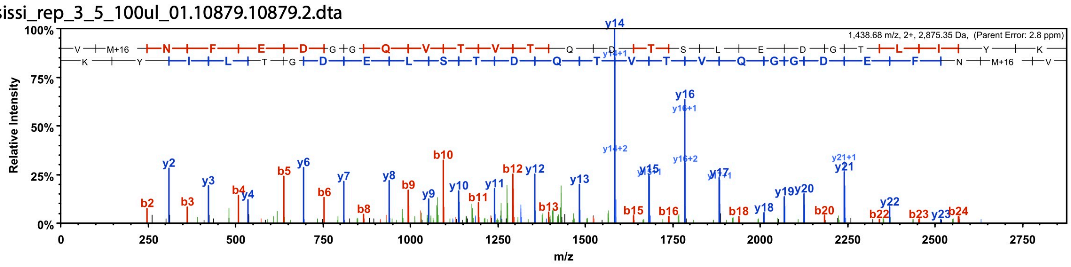


(R)VMNFEDGG**S**VTVTQDTSLEDGTLIYK(V)

s_s5_rep3_100ul_01.05091.05091.2.dta

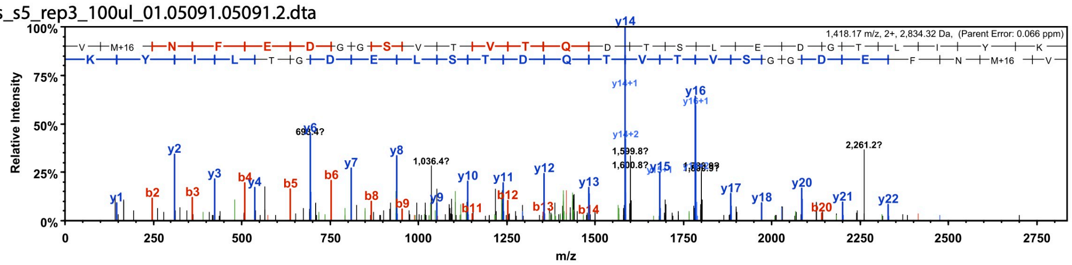


(R)VMNFEDGG**Y**VTVTQDTSLEDGTLIYK(V)

sissi_rep_3_5_100ul_01.12077.12077.2.dta

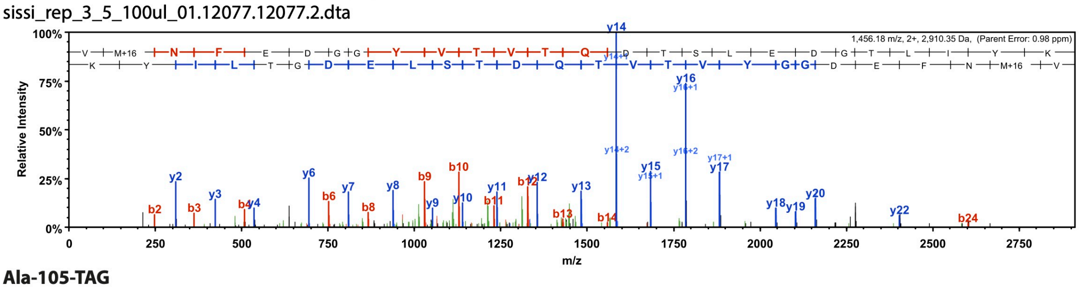


**Ala-105-TAG**

(R)VMNFEDGG**A**VTVTQDTSLEDGTLIYK(V)

sissi_rep_3_6_100ul_01.11857.11857.2.dta

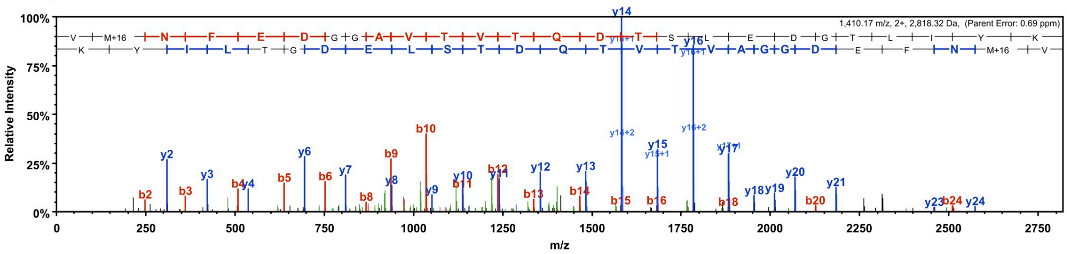

(R)VMNFEDGG**K**(V)
sissi_6_rep3_100ul.02714.02714.2.dta

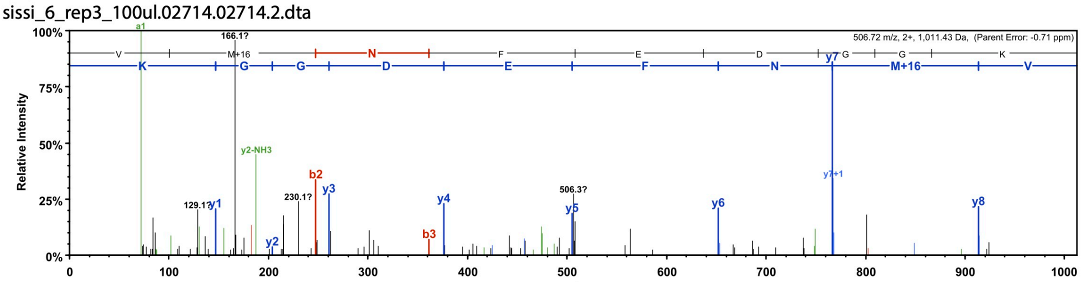


(R)VMNFEDGG**Q**VTVTQDTSLEDGTLIYK(V)
sissi_rep_3_6_100ul_01.13331.13331.2.dta

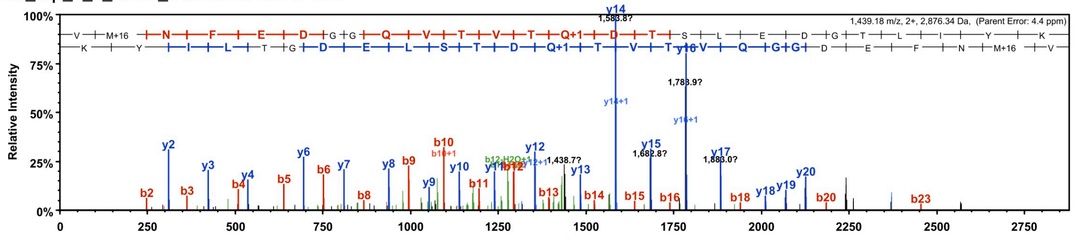


(R)VMNFEDGG**S**VTVTQDTSLEDGTLIYK(V)
s_s5_rep3_100ul_01.05091.05091.2.dta

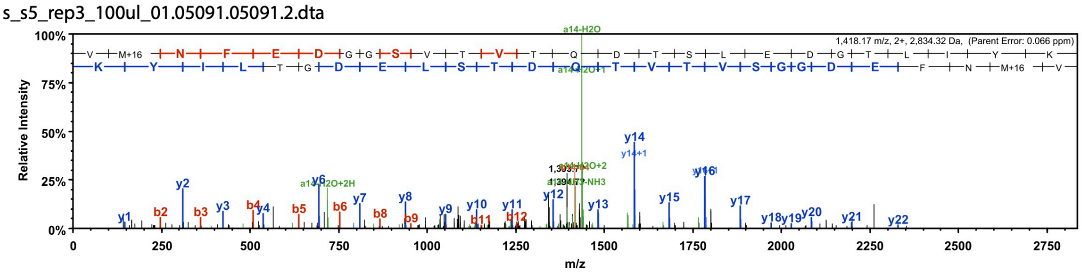


(R)VMNFEDGG**W**VTVTQDTSLEDGTLIYK(V)
s_s6_rep3_100ul_01.06082.06082.2.dta

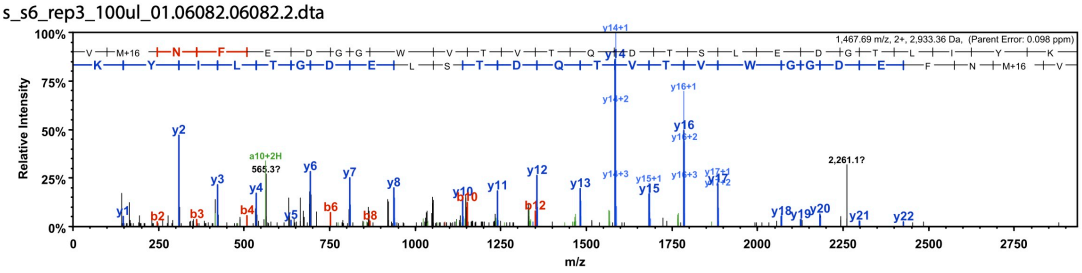


(R)VMNFEDGG**Y**VTVTQDTSLEDGTLIYK(V)
sissi_rep_3_6_100ul_01.12443.12443.2.dta

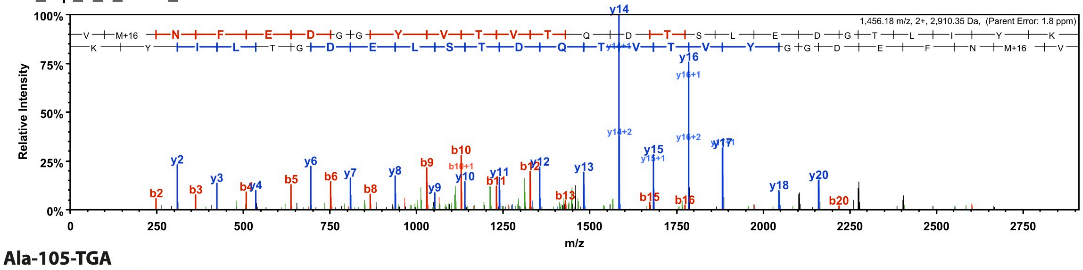


**Ala-105-TGA**
(R)VMNFEDGG**C**VTVTQDTSLEDGTLIYK(V)
s3_lane9_50ul.43349.43349.2.dta

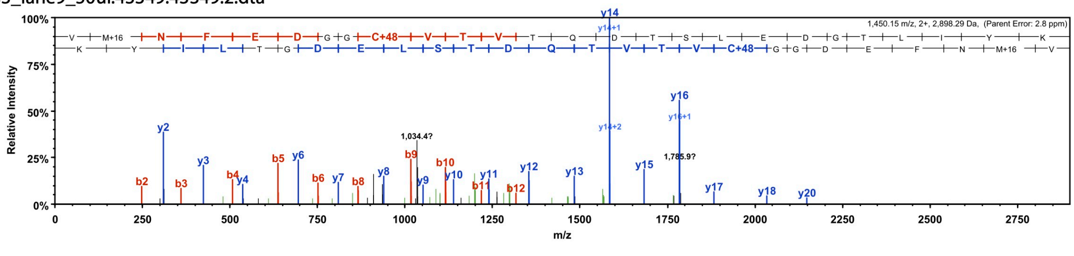

(R)VMNFEDGG**W**VTVTQDTSLEDGTLIYK(V)
s3_lane9_50ul.75236.75236.2.dta

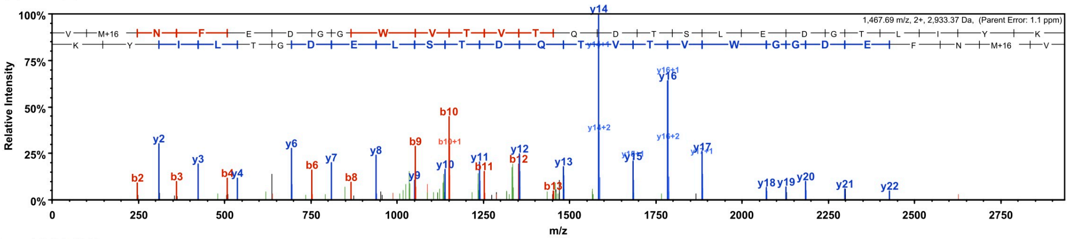


**Pro-135-TAG**

(K)LRGTNFPPDG**K**(V)
s2_lane3_11_50ul_210629211931.09651.09651.2.dta

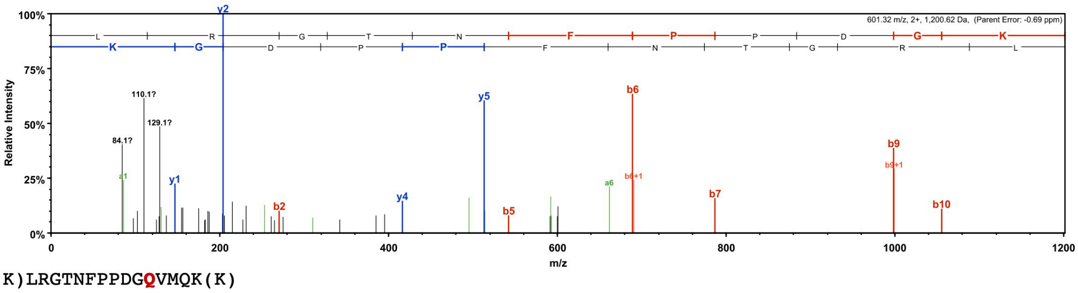


(K)LRGTNFPPDG**Q**VMQK(K)
s2_lane3_11_50ul_210629211931.14799.14799.2.dta

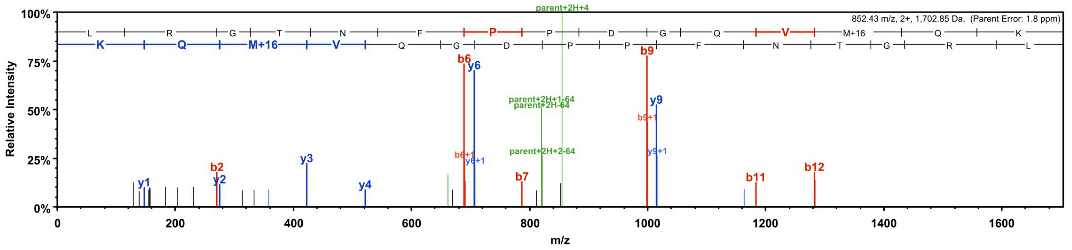


(K)LRGTNFPPDG**Y**VMQK(K)
s2_lane3_11_50ul_210629211931.20047.20047.2.dta

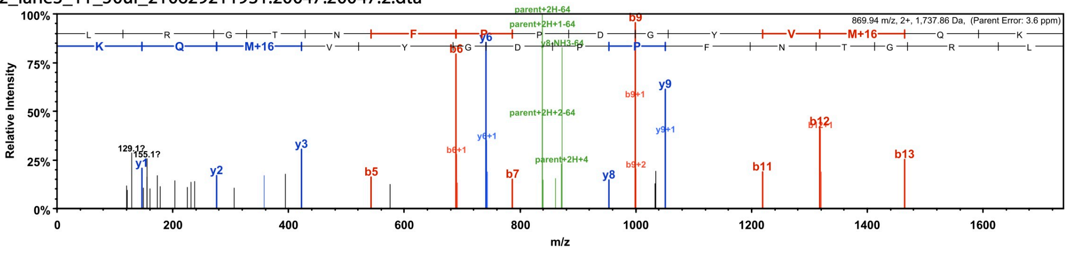


**Pro-135-TGA**

(K)LRGTNFPPDG**C**VMQK(K)
s1_lane9_50ul.15076.15076.2.dta

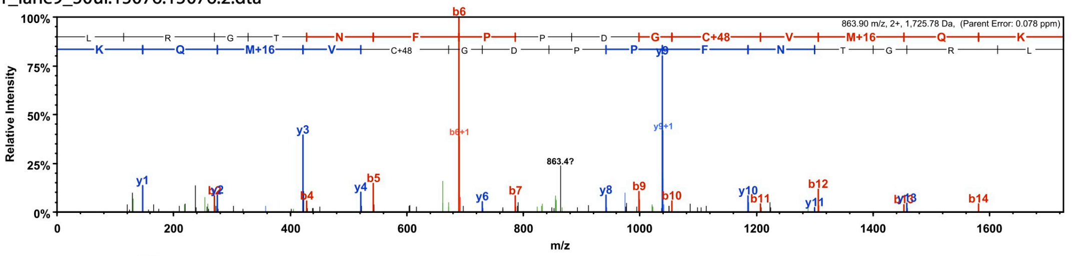


(K)LRGTNFPPDG**W**VMQK(K)
s1_lane9_50ul.18960.18960.2.dta

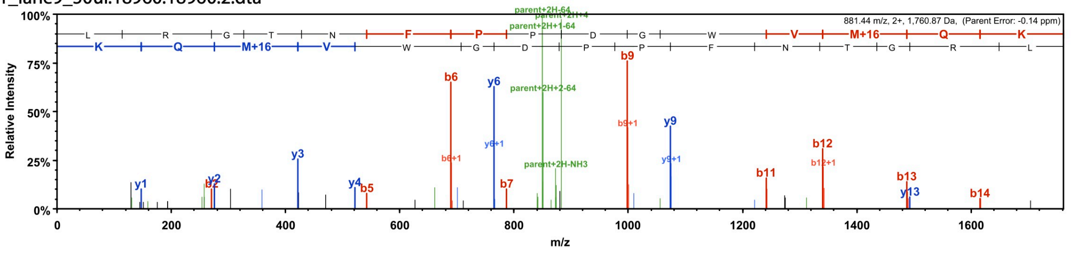

**Glu-145-TAG**

(K)TmGWQASTER(L)

s_8_2.03361.03361.2.dta

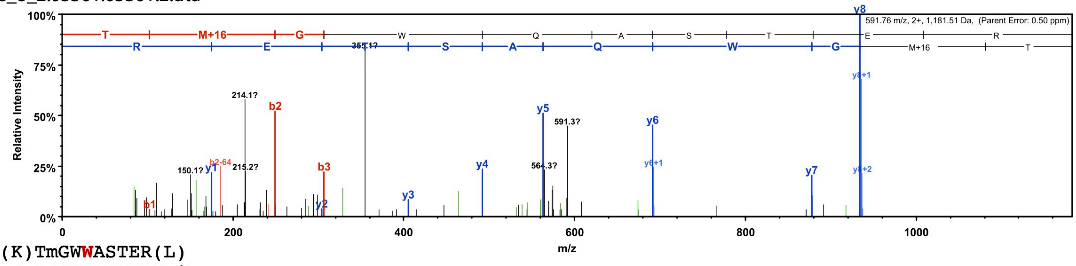


(K)TmGWWASTER(L)

s_8_2.06897.06897.2.dta

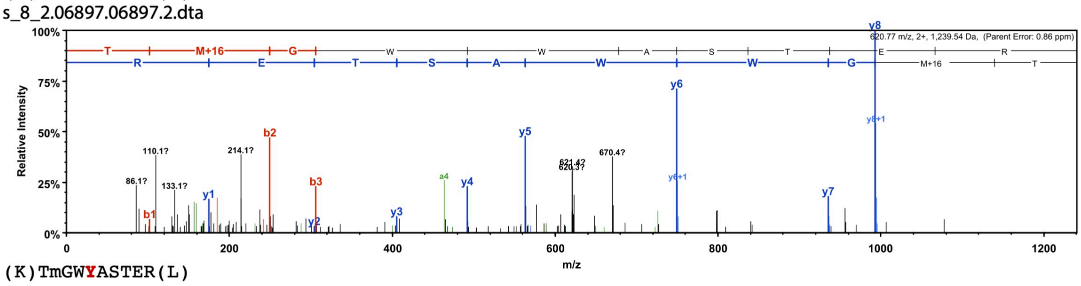


(K)TmGWYASTER(L)

s_8_2.04175.04175.2.dta

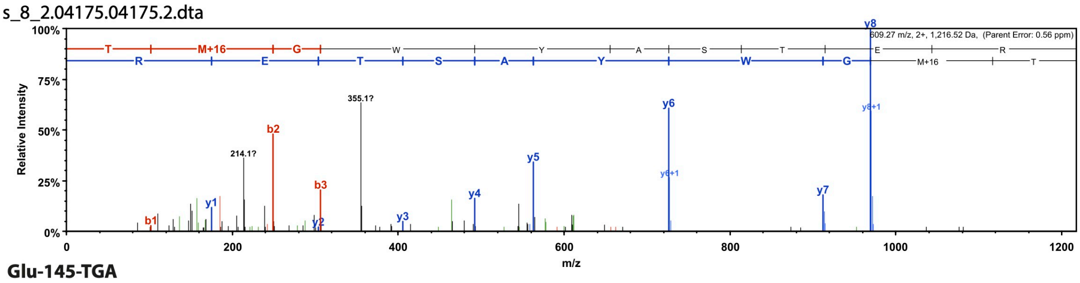


**Glu-145-TGA**

(K)TMGWEASTER(L)

s_s7_rep3_100ul_01.02210.02210.2.dta

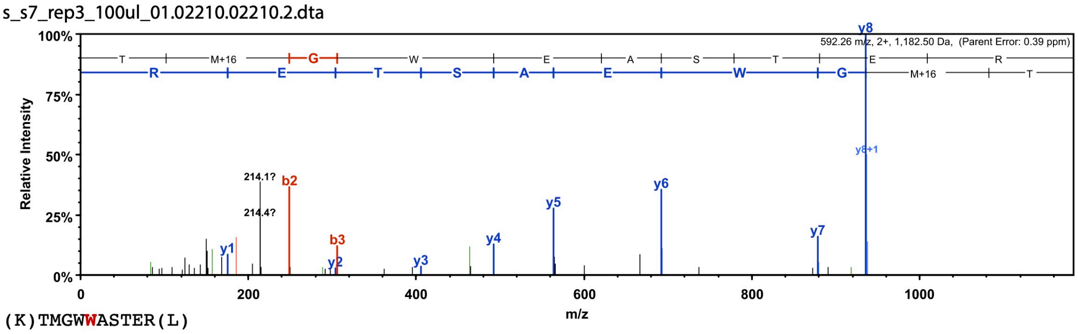


(K)TMGWWASTER(L)

sissi_7_rep3_100ul.12372.12372.2.dta

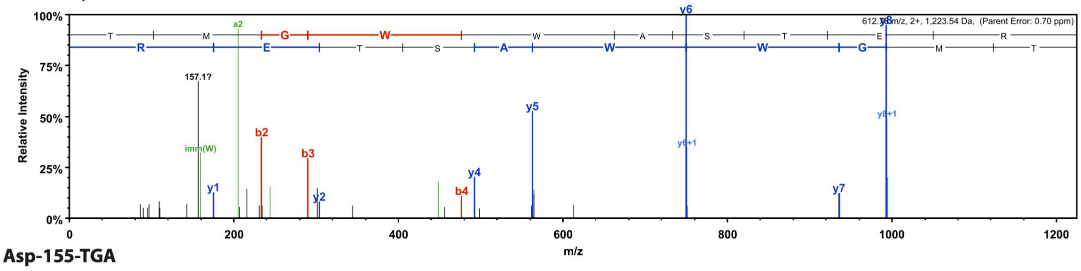


**Asp-155-TGA**

(R)LYPEDGVLK(G)

s_s4_2_100ul_02.02942.02942.2.dta

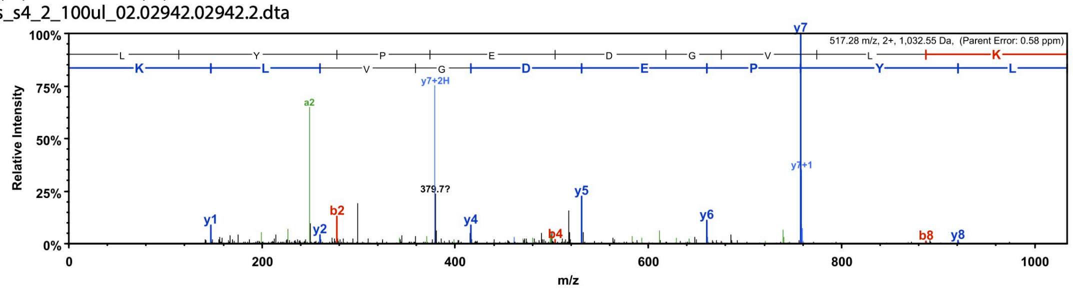

(R)LYPEWGVLK(G)

4_50ul_2.15909.15909.2.dta

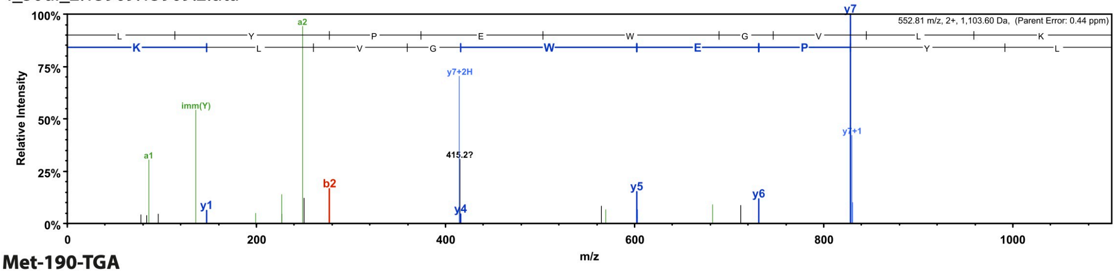


**Met-190-TGA**

(K)KPVQCPGAYNVDR(K)

m190_1_20220125103726.03059.03059.2.dta

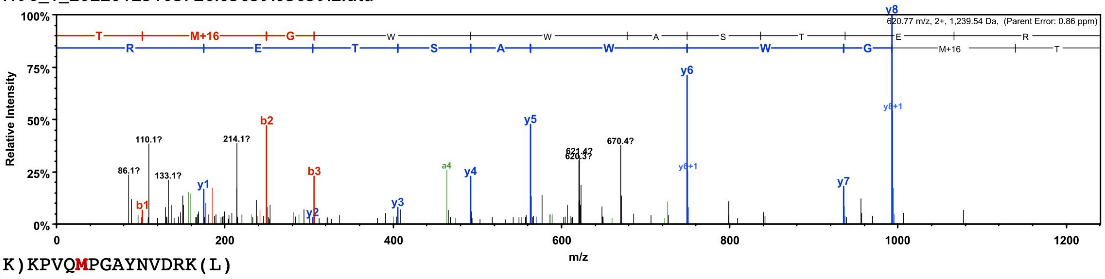


(K)KPVQMPGAYNVDRK(L)

s_m190_100ul_01.02587.02587.3.dta

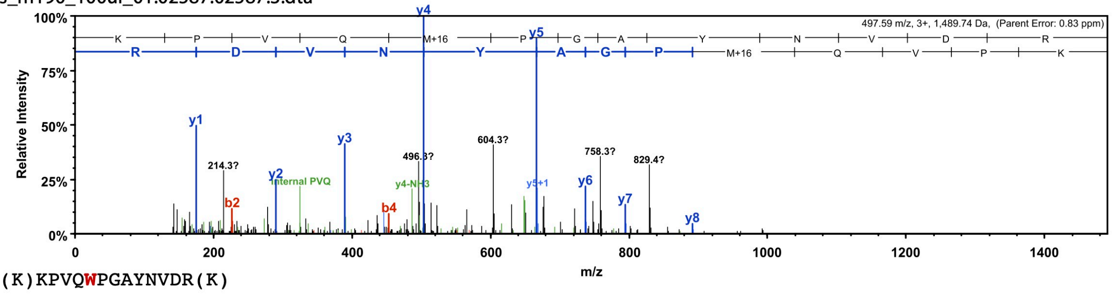


(K)KPVQWPGAYNVDR(K)

m190_1_20220125103726.06315.06315.2.dta

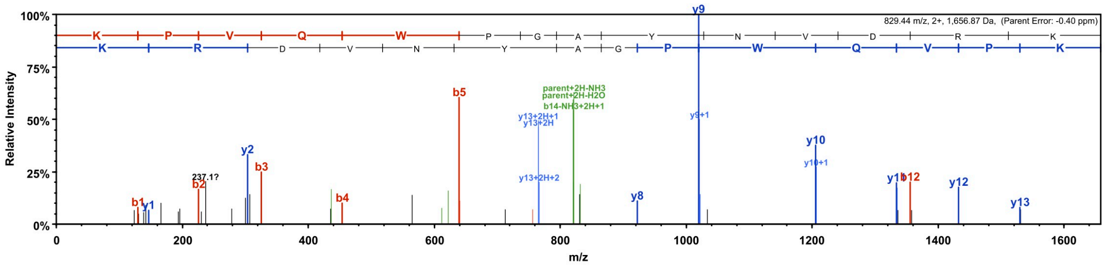


**Figure S6. Representative MS-spectra of reporters reveal the amino acids misincorporated at premature stop codon sites.** We selected 10 reporters for targeted mass spectrometry analysis. The spectra correspond to the peptides that cover the stop codon site. The amino acid that was inserted at the stop codon site is highlighted in red (see Supplementary Table S1and Method section for quantification of amino acid misincorporations).

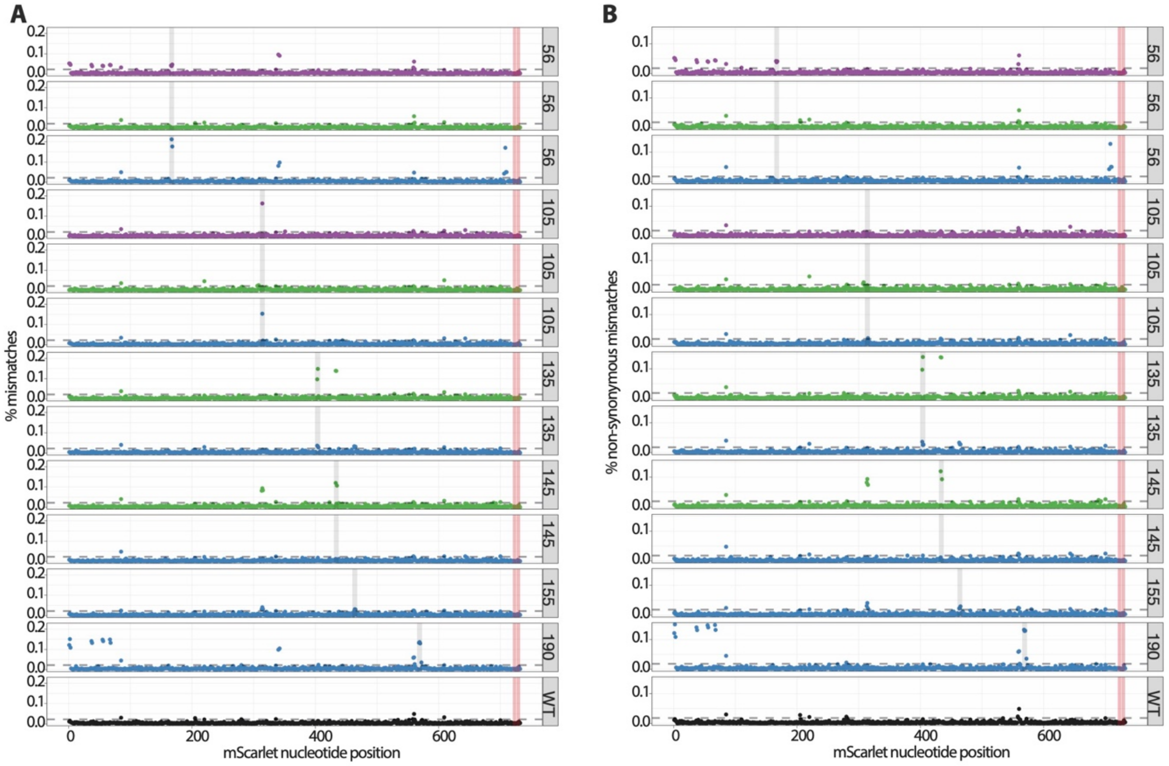


**Figure S7. RNA polymerase inaccurancy at premature stop codons revealed by RNA-seq data.** The percentages of nucleotide mismatches are shown along the mScarlet mRNA sequence for selected reporters. While most positions have a very low error rate, a few positions have a higher percentage of mismatches (A) and mismatches that result in an amino acid change (B, non-synonymous mismatches). The premature stop codon is indicated with grey shades and the canonical stop codon at the C-terminal is highlighted in red. There is no difference between reporters that harbor different stop codons, TAA (purple), TAG (green), TGA (blue). The WT sequence is shown in black.

#### ASWKPLLNLLP

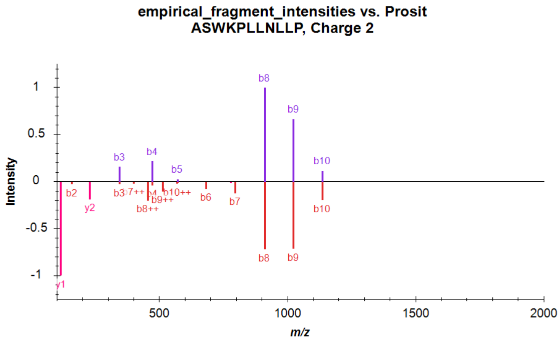


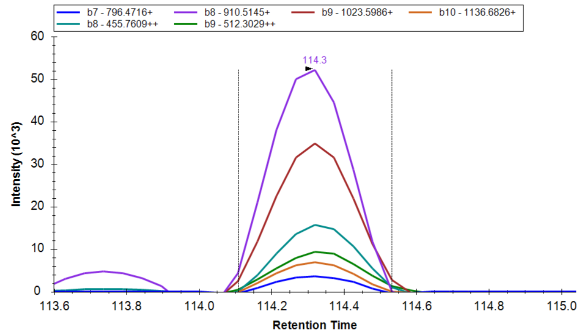


#### CGHSQQK

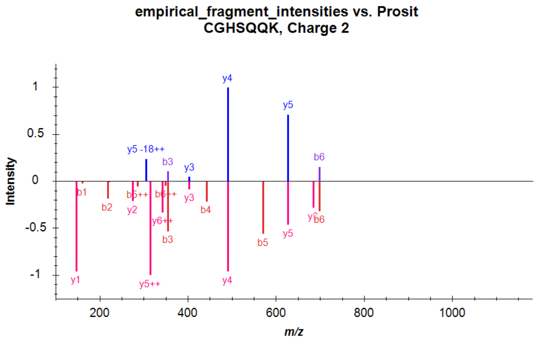


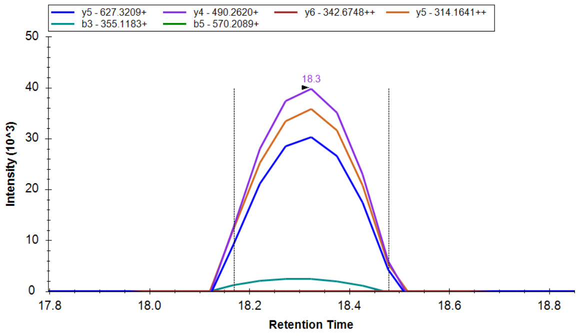


#### DDEAEKTEINGVAK

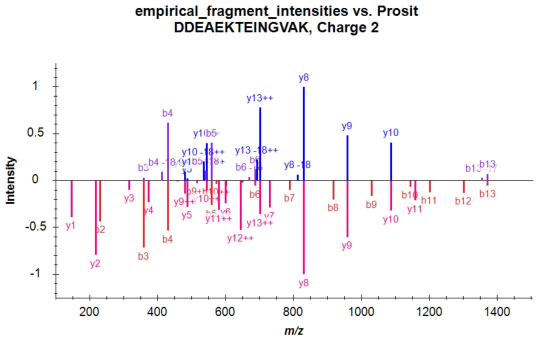


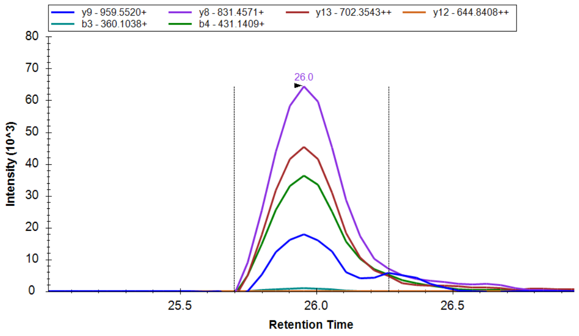


#### EITMGKTQPLPILITGGGR

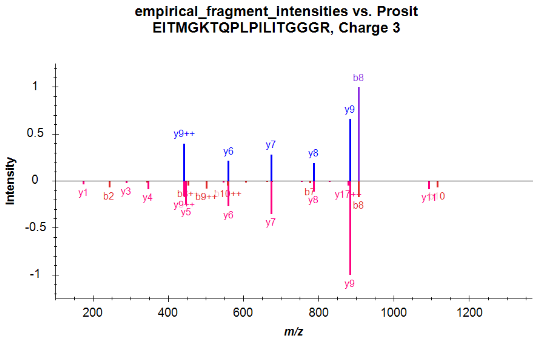


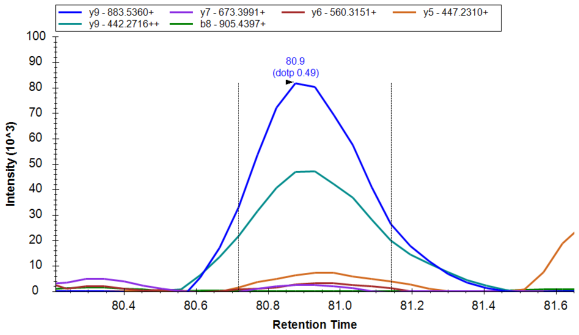

#### GPAAVNVTAIWSNPLI

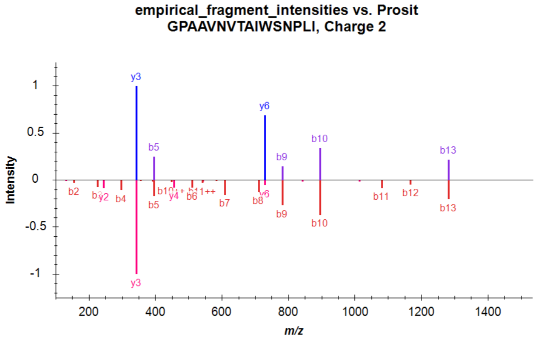


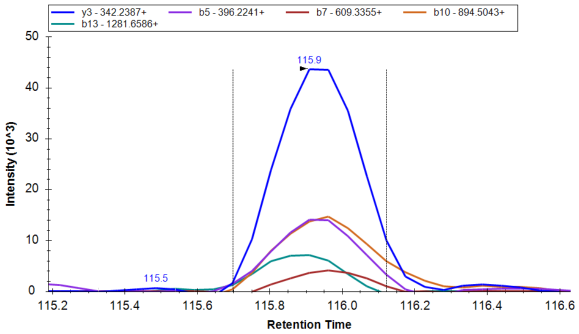


#### GSSWWSSVPLSDQMSR

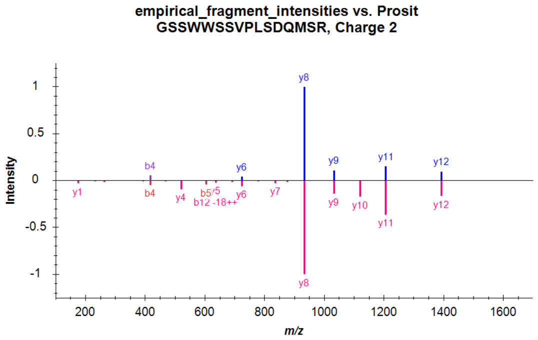


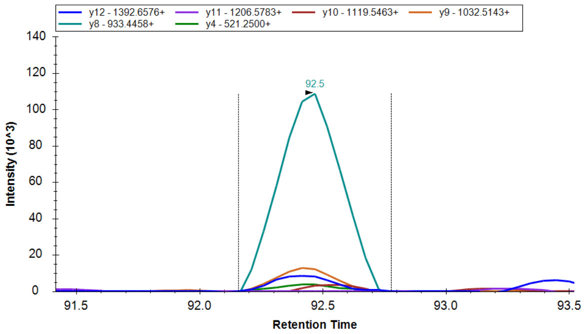


#### GVFAPLQIICVV

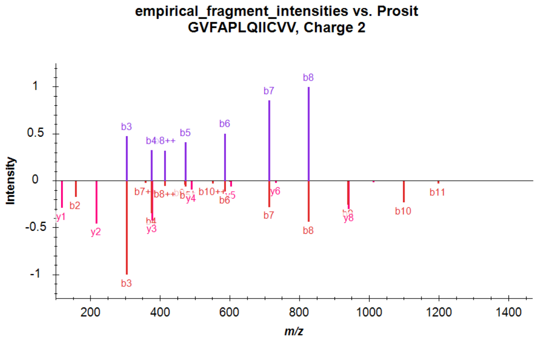


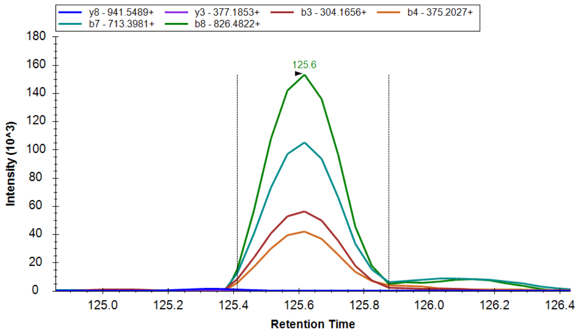


#### ILISFIR

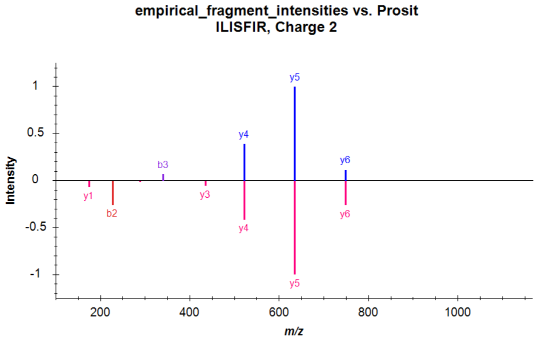


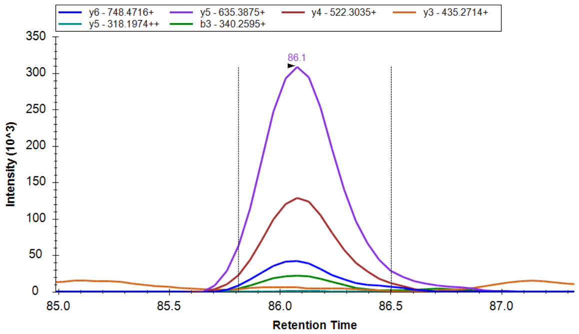

### KAGEAAVTVK

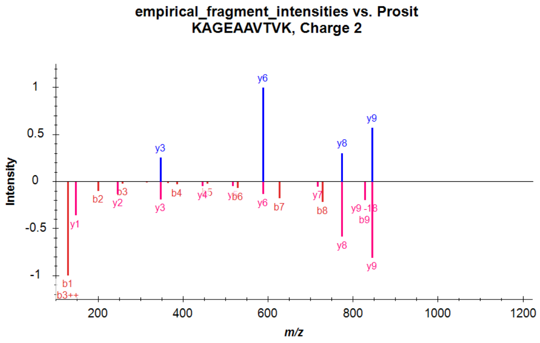


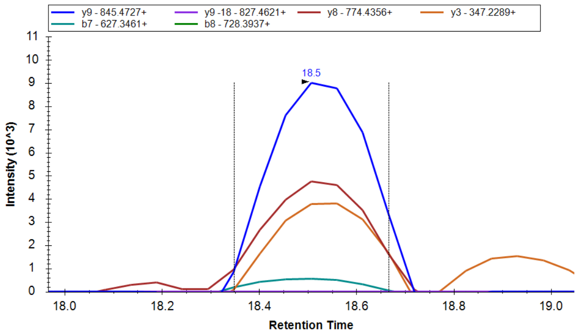


### KVAPGQNIASSR

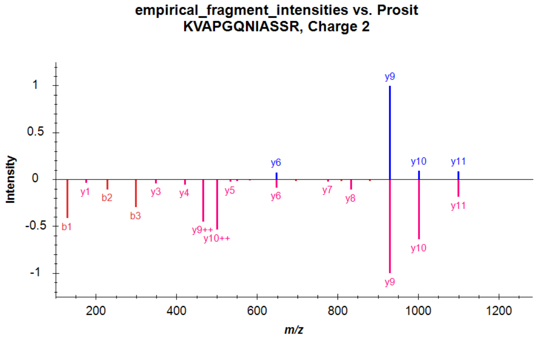


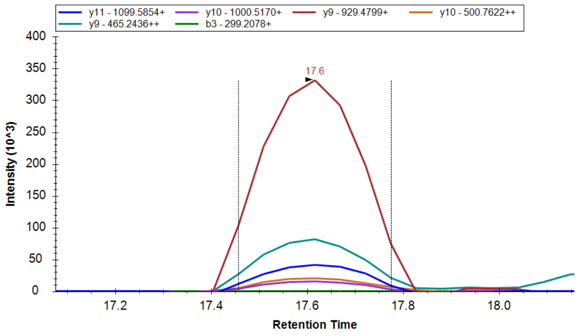


### LAHQAMTLK

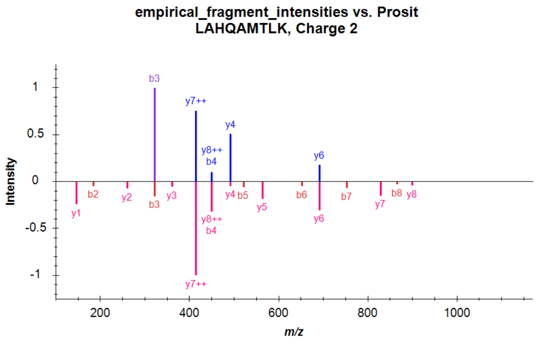


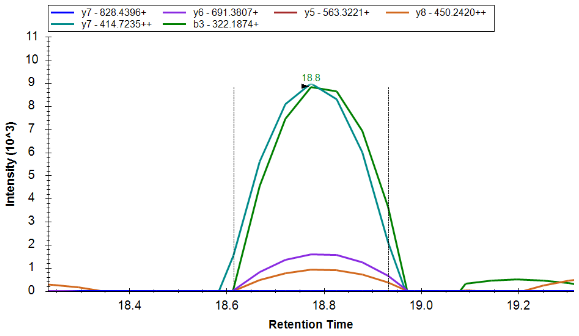


### NIAATLAIGMRNAGMQGR

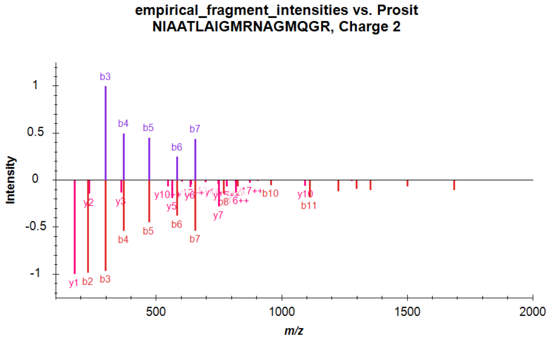


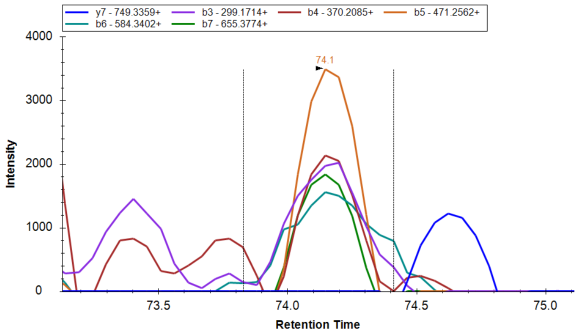

#### QPALGYLNCTPK

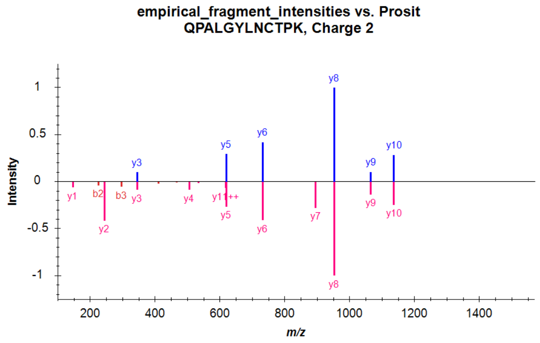


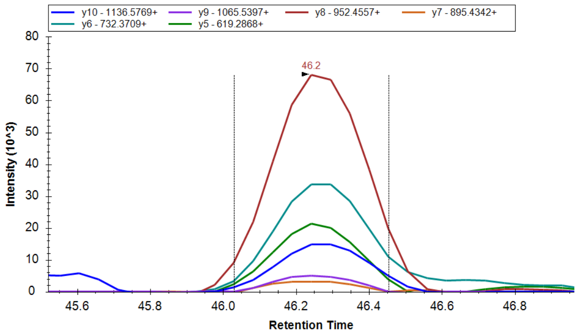


#### QPALGYLNWTPK

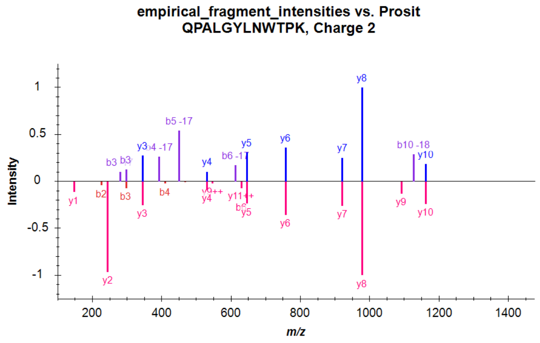


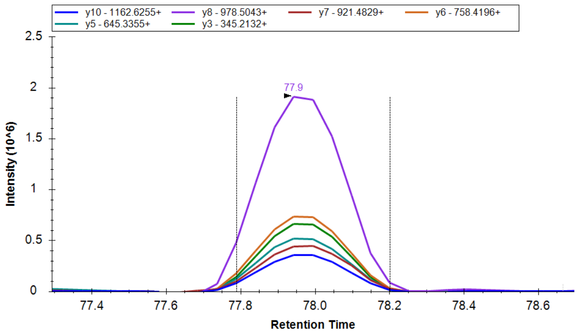


#### SEDAMSTQLDPTQLAIEFLR

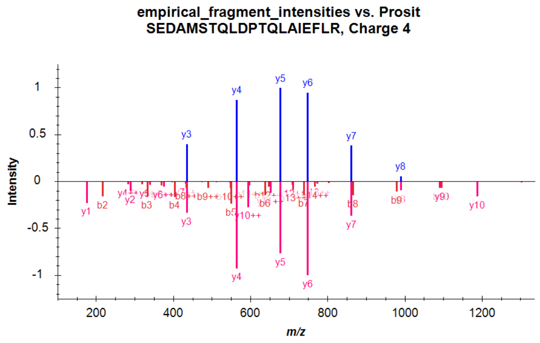


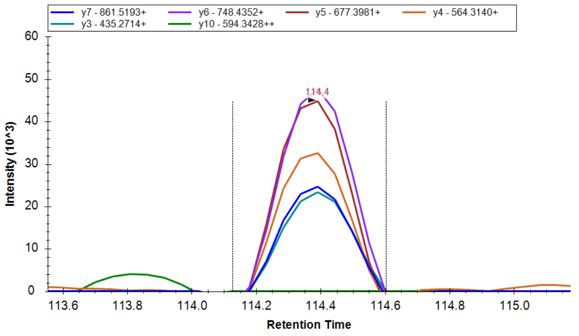


#### VLLIPWNR

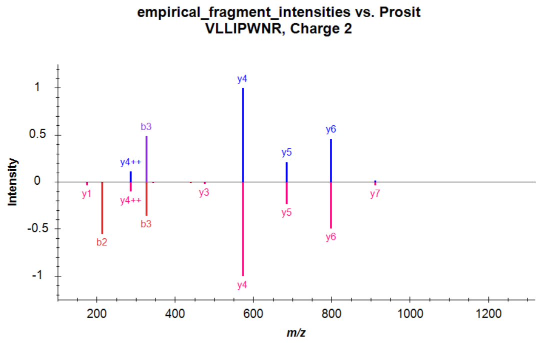


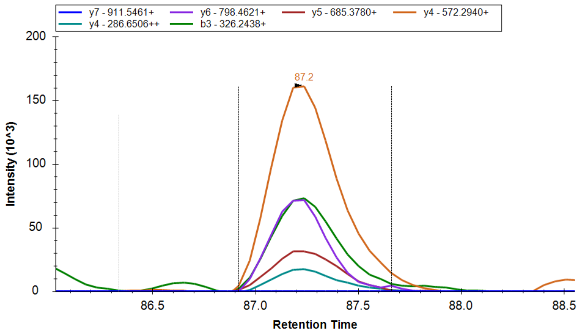

VYAGNEHNHAAQQPQVLDICSGL

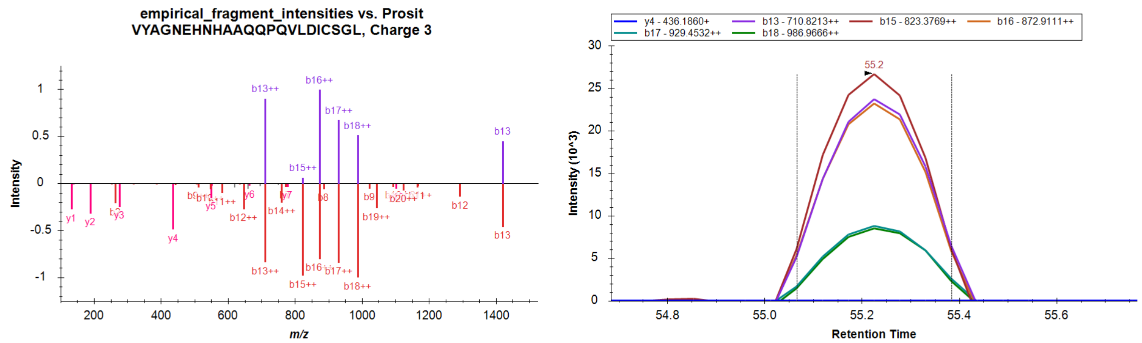


**Figure S8.** Evidence for peptides in Table S2 showing stop codon miscoding in the *E. coli* proteome. The data shown represent the evidence used by DIA-NN to identify the peptides. For each peptide, left panel shows a mirror plot of empirical relative fragment intensities extracted by DIA-NN as they appear in the output spectral library (upper spectrum), compared to Prosit prediction (lower spectrum; Prosit spectra predicted with NCE 31, taking into account the offset of 6 between Prosit and Orbitrap NCE[1]). Note that DIA-NN does not take into account the ions b1, b2, y1, and y2, even when predicted to be intense, due to a high likelihood of interference. For this reason, the dot product calculated by Skyline is not applicable and not shown. Right panels show chromatogram (with Savitsky-Golay smoothing) of fragments in the output spectral library from a single replicate, with peak boundaries as reported in the DIA-NN main output table.

**Data S1. Fluorescent readout of microscopy images (Fluorescence.zip).**
The readout of the image analysis are listed below for E.coli cells grown at different conditions. The files contain fluorescnet data for cells of all reporters (, WT mScaret (positive control, Control-WT) and two negative controls (Control-Col1 and Control-Col2).

- alldata_AHT.csv – AHT calibration data corresponding to Figure 1B.
- alldata_LB_18C.csv – fluorescent measuerements of cells grown at 18°C in LB corresponding to Figure S2A.
- alldata_LB_25C.csv – fluorescent measuerements of cells grown at 25°C in LB corresponding to Figure S2B.
- alldata_LB_37C.csv – fluorescent measuerements of cells grown at 37°C in LB corresponding to Figure S2C.
- alldata_LB_42C.csv – fluorescent measuerements of cells grown at 42°C in LB corresponding to Figure S2D.
- alldata_M9HCA_37C.csv - corresponding to Figure S4 pannel B.
- alldata_M9HG_37C.csv - corresponding to Figure S4 pannel A.
- alldata_M9_18C.csv – fluorescent measuerements of cells grown at 18°C in M9 corresponding to Figure S2E.
- alldata_M9_25C.csv – fluorescent measuerements of cells grown at 25°C in M9 corresponding to Figure S2F.
- alldata_M9_37C.csv – fluorescent measuerements of cells grown at 37°C in M9 corresponding to Figure S2G.
- alldata_M9_42C.csv – fluorescent measuerements of cells grown at 42°C in M9 corresponding to Figure S2A.

**Data S2. Sequencing data of the RNA-seq experiments (RNAseq.zip).**

- Ala105taa_bwa_sorted_only_mapped.bam
- Ala105tag_bwa_sorted_only_mapped.bam

- Ala105tga_bwa_sorted_only_mapped.bam
- Asp155tga_bwa_sorted_only_mapped.bam
- Glu145tag_bwa_sorted_only_mapped.bam
- Glu145tga_bwa_sorted_only_mapped.bam
- Met190tga_bwa_sorted_only_mapped.bam
- Pro056taa_bwa_sorted_only_mapped.bam
- Pro056tag_bwa_sorted_only_mapped.bam
- Pro056tga_bwa_sorted_only_mapped.bam
- Pro135tag_bwa_sorted_only_mapped.bam
- Pro135tga_bwa_sorted_only_mapped.bam
- Reference_to_plasmidASK_mScarlet.fa
- WT_bwa_sorted_only_mapped.bam

**Data S3. Sequencing data of the DNA-seq experiments (DNA-seq.zip).**

- Pro-56-taa-F-Premixed
- Pro-56-tag-F-Premixed
- Pro-56-tga-F-Premixed
- Ala-105-taa-F-Premixed.ab1
- Ala-105-tag-F-Premixed.ab1
- Ala-105-tga-F-Premixed.ab1
- Pro-135-tag-F-Premixed.ab1
- Pro-135-tga-F-Premixed.ab1
- Glu-145-tag-F-Premixed.ab1
- Glu-145-tga-F-Premixed.ab1
- Asp-155-tga-F-Premixed.ab1
- Met-190-tga-F-Premixed.ab1

**Table S1. Mass spectrometry analysis of the reporters revealed that stop codon miscoding primarily occur due to amino acid misincorporations.** TGA was almost always replaced by tryptophan (W) and, in lower frequency, by cysteine (C). glutamic acid (E), aspartic acid (D), and methionine (M). Misincorporation of glutamic acid (E), aspartic acid (D), and methionine (M), at the TAG position was a minor process. TAA and TAG were replaced by glutamine (Q), tyrosine (Y), and lysine (K), where all three amino acids had comparable misincorporation rates. Misincorporation of alanine (A) and serine (S), at the TAA position and alanine (A), serine (S), and tryptophan (W), at the TAG position was a minor process.

| aa position mutated to stop codon | stop codon | peptides detected by ms[1] | misincorporated amino acids detected by ms | relative abundance (%)[2] |
|---|---|---|---|---|
| 105 | TAA | (R) VMNFEDGG**X**VTVTQDTSLEDGTLIYK (V)<br>(R) VMNFEDGG**K** | Q K **Y** A S | Q(58), Y(15), A(26)<br>S(<1), K |
| 105 | TAG | (R) VMNFEDGG**X**VTVTQDTSLEDGTLIYK (V)<br>(R) VMNFEDGG**K** | Q K Y A S W | Q(77), Y(14), A(9)<br>S(<1), W(<1), K |
| 105 | TGA | (R) VMNFEDGG**X**VTVTQDTSLEDGTLIYK (V) | W C | W(99), C(1) |
| 135 | TAG | (K) LRGTNFPPDG**X**VMQK (K)<br>(R) GTNFPPDG**X**VMQK (K)<br>(K) LRGTNFPPDG**K** (V)<br>(R) GTNFPPDG**K** (V) | Q Y K | Q(73), Y(27)<br>K |
| 135 | TGA | (K) LRGTNFPPDG**X**VMQK (K)<br>(R) GTNFPPDG**X**VMQK (K) | W C | W(99), C(1) |
| 145 | TAG | (K) TMGW**X**ASTER (L)<br>(K) KTMGW**X**ASTER (L) | Q Y W | Q(39), Y(58), W(2) |
| 145 | TGA | (K) TMG**X**ASTER (L)<br>(K) KTMGW**X**ASTER (L) | W E | W(>99)<br>E(<1) |
| 155 | TGA | (R) LYPE**X**GVLK (G)<br>(R) LYPE**X**GVLKGDIK (M) | W D | W(>99)<br>D(<1) |
| 190 | TGA | (K) KPVQ**X**PGAYNVDR (K)<br>(K) AKKPVQ**X**PGAYNVDR (K) | W C M | W(99), C(~1)<br>M(<1) |

[1] - Only peptides covering mutated position are shown; **X** or **K** (for Lys) designate incorporated amino acid

[2] - Calculated as described in Materials and Methods for the corresponding forms of the peptide comprising mutated position; K was not included.

**Table S2. Proteome-wide mass spectrometry reveals the expression of non-coding sequences by stop codon miscoding events in *E. coli*.** Peptides indicating SCM events are listed with modification and charge state as reported by DIA-NN. TGA was more error-prone stop codon than TAA. We did not detect any evidences of SCM in TAG proteins, probable due to is low representation in the *E.coli* genome (8%). We detected more cases of SCM in *E.coli* samples grown at l8°C than at 37°C (14 vs. 11, see also Fig 2A and C).

| Peptide[1] | Gene | Type | Stop Codon | Canonical Protein Length (aa) | Peptide Start | Peptide End | Inserted aa | IDs 18°C | IDs 37°C | Found at 18°C[2] | Found at 37°C[2] | Median log2 Int 18°C[3] | Median log2 Int 37°C[3] | Log Change | P-value[4] | Significant[5] |
|---|---|---|---|---|---|---|---|---|---|---|---|---|---|---|---|---|
| (R)GSS**W**WSSVPLSDQMSR(R) | *astC* | Covers_Stop | TGA | 406 | 403 | 418 | W | 6 | 0 | TRUE | FALSE | 19.5398 | | | 1 | FALSE |
| (R)GVFAPLQIICVV(*) | *birA* | Past_Stop | TAA | 321 | 324 | 335 | X | 4 | 6 | TRUE | TRUE | 16.9271 | 18.3338 | -1.4067 | 0.0537 | FALSE |
| (K)GPAAVNVTAI**W**SNPLI(*) | *cspC* | Covers_Stop | TGA | 69 | 59 | 74 | W | 5 | 0 | TRUE | FALSE | 18.3445 | | | 1 | FALSE |
| (R)**C**GHSQQK(Y) | *dcm* | Covers_Stop | TAA | 472 | 472 | 478 | C | 5 | 4 | TRUE | TRUE | 18.4527 | 19.1635 | -0.7108 | 0.4073 | FALSE |
| (R)NIAATLAIGMRNAG**M**QGR(A) | *garK* | Covers_Stop | TGA | 381 | 367 | 384 | M | 3 | 6 | TRUE | TRUE | 15.5465 | 17.0081 | -1.4616 | 0.0006 | TRUE |
| (K)VLLIP**W**NR(G) | *gatD* | Covers_Stop | TGA | 346 | 341 | 348 | W | 6 | 5 | TRUE | TRUE | 22.0872 | 19.6054 | 2.4818 | 0 | TRUE |
| (K)SEDAMSTQLDPTQLAIEFLR(R) | *menI* | Past_Stop | TGA | 136 | 166 | 185 | X | 2 | 3 | FALSE | TRUE | 17.3435 | 16.1639 | 1.1796 | 0.6136 | FALSE |
| (R)KAGEAA**V**TVK(N) | *mhpF* | Covers_Stop | TGA | 316 | 310 | 319 | V | 5 | 3 | TRUE | TRUE | 16.2749 | 17.9175 | -1.6426 | 0.0079 | FALSE |
| (K)AS**W**KPLLNLLP(*) | *ribB* | Covers_Stop | TGA | 217 | 215 | 225 | W | 6 | 6 | TRUE | TRUE | 19.8146 | 18.1035 | 1.7111 | 0 | TRUE |
| (K)VYAGNEHNHAAQQPQVLDI**C**SGL(*) | *rplM* | Covers_Stop | TAA | 142 | 123 | 145 | C | 4 | 5 | TRUE | TRUE | 18.6309 | 16.5107 | 2.1202 | 0.0102 | FALSE |
| (K)QPALGYLN**C**TPK(R) | *rpsG* | Covers_Stop | TGA | 179 | 171 | 182 | C | 6 | 6 | TRUE | TRUE | 18.9541 | 18.8967 | 0.0574 | 0.6483 | FALSE |
| (K)QPALGYLN**W**TPK(R) | *rpsG* | Covers_Stop | TGA | 179 | 171 | 182 | W | 6 | 6 | TRUE | TRUE | 44.146 | 31.6105 | 12.5355 | 0.0119 | FALSE |
| (R)DDEAEK**T**EINGVAK(C) | *ybaM* | Covers_Stop | TGA | 53 | 47 | 60 | T | 1 | 4 | FALSE | TRUE | 20.9835 | 20.7976 | 0.1859 | 1 | FALSE |
| (K)KVA**P**GQNIASSR(R) | *ybaQ* | Covers_Stop | TAA | 113 | 110 | 121 | P | 3 | 2 | TRUE | FALSE | 20.6712 | 19.0116 | 1.6596 | 0.958 | FALSE |
| (K)EITMGKTQPLPILITGGGR(R) | *ydgI* | Past_Stop | TAA | 460 | 470 | 488 | X | 3 | 1 | TRUE | FALSE | 20.6283 | 18.9162 | 1.7121 | 1 | FALSE |
| (K)LAHQAMTLK(L) | *yeiI* | Past_Stop | TAA | 362 | 391 | 399 | X | 4 | 0 | TRUE | FALSE | 16.791 | | | 1 | FALSE |
| (K)ILIS**F**IR(K) | *yrhD* | Covers_Stop | TAA | 51 | 47 | 53 | F | 4 | 5 | TRUE | TRUE | 22.0012 | 21.6268 | 0.3744 | 0.189 | FALSE |

[1] amino acids inserted at the Stop codon are **bold**, underlined residues are modified: carbamidomethylation (+57) at C, oxidation (+18) at M; previous and following amino acids indicated in brackets, * indicates end of sequence

[2] identification in a given condition (18 or 37ºC) is defined as detection in at least 3/6 replicates of that condition
[3] median of log2 intensity values as reported by DIA-NN over replicates of the condition; if empty, peptide was not detected in the condition
[4] unadjusted p-value of two-sided Student's t-test on log2-transformed intensities of the two temperature conditions
[5] after adjustment by Benjamini-Hochberg method
